# Macrophage migration is differentially regulated by distinct ECM components

**DOI:** 10.1101/2023.04.27.538597

**Authors:** Matthew W. Stinson, Alexander J. Laurenson, Jeremy D. Rotty

**Affiliations:** Uniformed Services University of the Health Sciences, Department of Biochemistry, Bethesda, MD, USA, 20814; The Henry M. Jackson Foundation for the advancement of Military Medicine, Bethesda, MD, USA, 20817.

**Author notes:** Corresponding author: Jeremy D. Rotty, Assistant Professor of Biochemistry, 4301 Jones Bridge Rd. Bethesda, MD 20814.

**Keywords:** Extracellular matrix, cell migration, myosin, ROCK, fibronectin, laminin, macrophage

## Abstract

Macrophages are indispensable for proper immune surveillance and inflammatory regulation. They also exhibit dramatic phenotypic plasticity and are highly responsive to their local microenvironment, which includes the extracellular matrix (ECM). The present work demonstrates that two fibrous ECM glycoproteins, fibronectin (FN) and laminin (LAM), elicit distinct morphological and migratory responses to macrophages in 2D environments. Laminin 111 inhibits macrophage cell spreading, but drives them to migrate rapidly and less persistently compared to cells on fibronectin. Differential integrin engagement and ROCK/myosin II organization helps explain why macrophages alter their morphology and migration character on these two ECM components. The present study also demonstrates that laminin 111 exerts a suppressive effect toward fibronectin, as macrophages plated on a LAM/FN mixture adopt a morphology and migratory character almost identical to LAM alone. This suggests that distinct responses can be initiated downstream of receptor-ECM engagement, and that one component of the microenvironment may affect the cell’s ability to sense another. Overall, macrophages appear intrinsically poised to rapidly switch between distinct migratory modes based on their ECM environments. The role of ECM composition in dictating motile and inflammatory responses in 3D and *in vivo* contexts warrants further study.

## INTRODUCTION

Macrophages are innate immune cells that function in host defense, antigen presentation, and tissue repair.^[1]^ Macrophages are derived from circulating bone-marrow monocytes, the fetal liver, or from yolk sac-progenitors.^[2]^ Despite the importance of macrophages in host defense and tissue repair, it is not fully understood how each component of the local microenvironment contributes to the vast array of macrophage inflammatory phenotypes and functions.^[3–5]^ It is well appreciated that macrophages are highly responsive to their environment. A single, well-studied example of this is their activation by cytokines and chemokines present in the microenvironment. These cues can shift over time, leading macrophages to adopt an array of inflammatory or anti-inflammatory profiles during physiological responses to damage or infection. In addition, there are many disease states that manifest alongside chronic inflammation, which may be due in part to dysregulated macrophage polarization.^[6–10]^ A better understanding of how the surrounding microenvironment elicits distinct macrophage functions may give an additional avenue to target dysregulated macrophage activation during pathological processes.

The extracellular matrix (ECM) is one such environmental cue that, compared to chemokines and cytokines, has been understudied from an immunomodulatory perspective. It has long been known that macrophages can be primed by substrate adhesion^[11, 12]^, and that fibronectin possesses an immunosuppressive character while collagen promotes TNF-α and CSF induction^[13]^. However, isoforms of fibronectin containing ‘extra domains’ (ED) have been implicated as pro-inflammatory via their ability to bind TLR4 ^[14, 15]^. Vitronectin has more recently been found to stimulate IL-6 and LIF secretion^[16]^. One especially compelling study noted that fibrinogen seems to potentiate LPS/IFNγ-stimulated inflammation, while fibrin seems to block it ^[17]^. Laminin induces MAPK signaling ^[18]^ and was one of several ECMs reported to induce pro-inflammatory cytokines as well ^[19]^. Both tissue resident macrophages and monocyte-derived populations at steady state or responding to infection or trauma encounter an array of ECM cues that can be tissue specific and time-dependent. Specific regions of the same tissue may even have dramatically different ECM compositions. An example of this is the laminin- and collagen IV-rich basement membrane compared to the underlying fibronectin- and collagen-rich connective tissue (which also can contain vitronectin).^[20]^ It is clear from the literature that these different ECM microenivronments are capable of differentially tuning macrophage behavior, but there is still much to learn about the molecular mechanisms that drive these responses.

Macrophages are highly responsive migratory cells that migrate through dense, 3-D extracellular matrices of varying composition to successfully resolve inflammation or address infection.^[21]^ The exact protein composition of an extracellular matrix is highly complex and mutable, as ECMs are tissue dependent and can fluctuate over time, especially during physiological processes like wound healing^[22–24]^. Despite the variety of fibrous ECM components, leukocyte cell migration studies tend to utilize fibronectin because fibronectin is secreted by macrophages, known to attenuate inflammatory signaling, and it induces spreading and adhesion.^[25–28]^ Macrophage anti-inflammatory behavior can also be enhanced by altering cell shape and size via mechanical means such as micropatterning, altering substrate rigidity or adjusting surface topography.^[29–31]^ Thus, the components of the extracellular matrix, and the effects they have on cell shape, size, and motility in the tissue, may influence local inflammatory signaling in a tissue-specific or region-specific manner.

Integrins are major ECM-sensing heterodimeric transmembrane receptors that are ubiquitously expressed. Each heterodimer has an affinity for one or more specific ECM components. Non-covalent interactions between α-(18 types) and β-(8 types) subunits form a total of at least 24 cell- and tissue-type dependent integrins, which are synthesized and heterodimerize in the endoplasmic reticulum and Golgi before they are transported to the cell surface.^[32, 33]^ Integrins must be in their active, unfolded conformation in order to interact with the microenvironment. Active integrins must be stimulated either by intracellular signals, such as talin and kindlin binding to the cytoplasmic tail (called “inside-out” signaling), and/or by mechanical forces induced by the ECM itself (i.e., strong adhesion pulling open partially active integrins, (called “outside-in” signaling). These signals can compete for access to adaptor and regulatory proteins, making it challenging to fully map out their signaling pathways. A greater understanding of how these convergent and divergent signaling pathways respond to specific microenvironmental components is required in order to better understand physiological processes that occur in complex, mutable 3-D environments like those found *in vivo*.

We have determined that the ECM components fibronectin and laminin can intrinsically alter the motility, morphology, and adhesion of macrophages through differential integrin activation. One the one hand, fibronectin induces slow, relatively persistent migration, as well as spreading, elongation, and strong adhesion by stimulating integrin pairs like αvβ1, αvβ3 and α5β1 that bind to fibronectin’s Arginine-Glycine-Aspartate (RGD). Conversely, laminin, signaling through α6β1 integrin, induces rapid, relatively random, slingshot-like migration, a balled-up morphology, and relatively weak adhesion. Myosin II and ROCK both contribute to the laminin phenotype. Whereas myosin II contributes to the high velocity and dynamic cell shapes changes induced by laminin, myosin-independent ROCK activity causes macrophages to adopt a balled-up morphology and to migrate less persistently than cells on fibronectin. Macrophages on fibronectin require integrins α_V_β_3_ and α_V_β_5_ to induce spreading and the relatively slower, more persistent fibronectin motility phenotype. Taken together, these data suggest that local ECM context alone dramatically influences macrophage cell morphology and migration by signaling through specific integrins to distinct actin-regulating pathways that allow macrophages to immediately and dynamically respond to their environments. Since ECM components like fibronectin and laminin can elicit changes in inflammatory and migratory signaling it is necessary to take these factors into account when conducting studies on immune cells *in vitro*. Furthermore, diseases that alter ECM secretion, localization, and organization could have detrimental or beneficial effects on macrophage inflammatory polarization when combined with cytokines secreted in the local tissue environment.

## RESULTS

### Macrophage cell morphology and motility are tuned by the extracellular matrix

We first plated macrophages on equal concentrations of various ECMs to test the hypothesis that the matrix microenvironment influences cell morphology and cytoskeletal organization. Macrophages produced F-actin at high levels and spread well in many ECM contexts (Fig. 1A), except when exposed to laminin 111 (hereafter referred to as ‘laminin’ or LAM). Laminin uniquely impaired cell spreading, caused macrophages to adopt a less elongated morphology, and suppressed F-actin polymerization (Fig. 1A-D). Conversely, fibronectin (FN) strongly stimulated cell spreading compared to the other ECMs (Fig. 1A, 1B), as well as supporting cellular elongation and F-actin polymerization (Fig. 1C, 1D). Statistical relationships between all ECMs tested can be found in Supplemental Figure 1. These data indicate that the local ECM environment is sufficient to alter cell morphology and the actin cytoskeleton.

**Figure 1:**
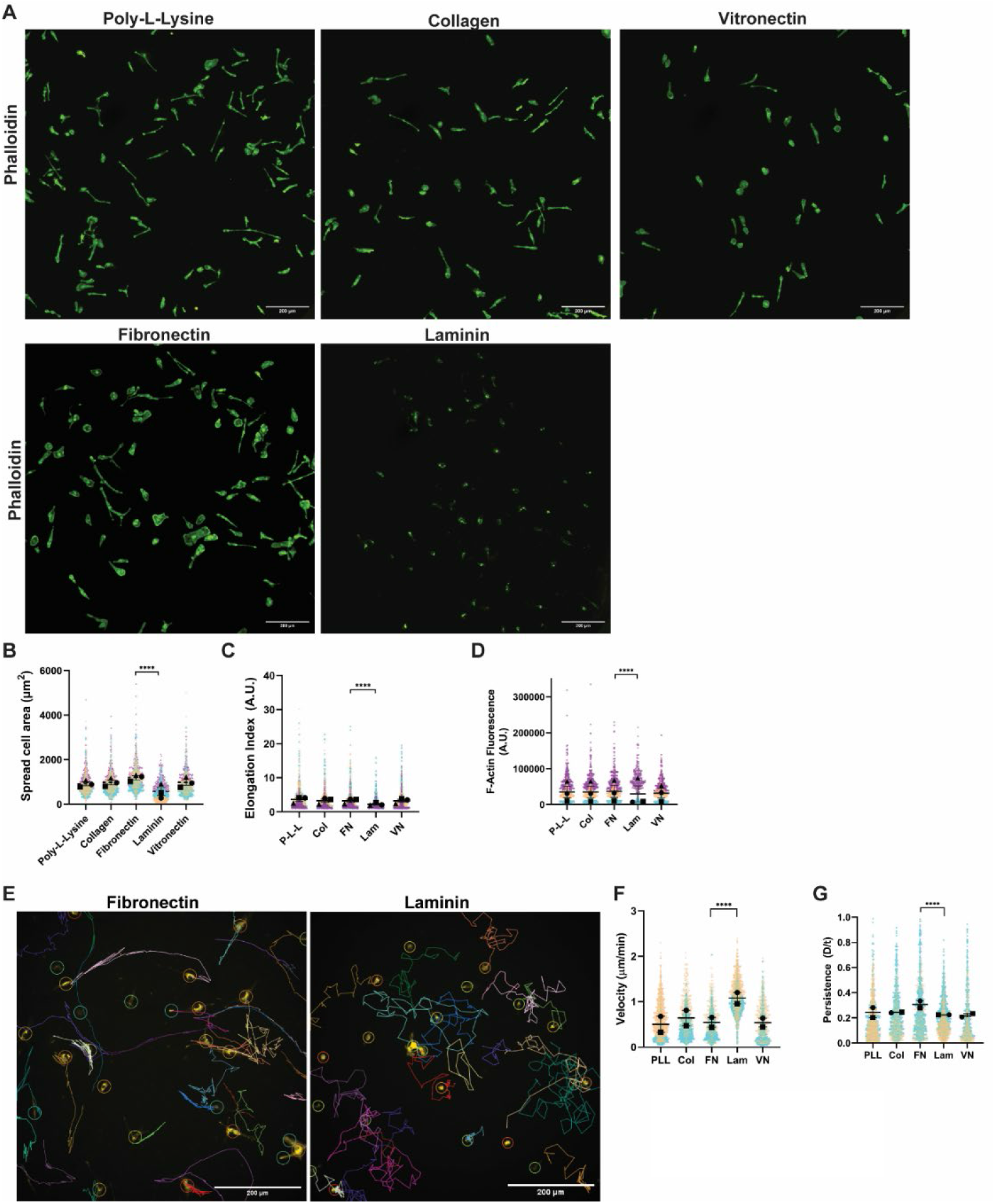
Macrophage cell morphology and motility are tuned by the extracellular matrix. (A) Representative images of fixed macrophages stained with Phalloidin 488 on 10 μg/mL Poly-L-Lysine (P-L-L), Collagen (Col), Fibronectin (FN), Laminin (Lam), or Vitronectin (VN). Scale bar = 200 microns in each image. (B) Quantification of spread cell area in microns squared (**** = p<0.0001, FN n=967 cells, Lam n=846 cells pooled from 3 independent experiments). Means and standard error of the mean for each experiment are represented with black symbols, all data points are plotted and each experimental run is color-coded. Statistical analysis was Kruskal-Wallis with Dunn multiple comparisons test. (C) Quantification of cellular elongation (A.U.). Greater elongation corresponds to a higher value in this analysis. (**** = p<0.0001, FN n=967 cells, Lam n=846 cells pooled from the same 3 independent experiments used to generate data in Figure 1B). Means and standard error of the mean for each experiment are represented with black symbols, all data points are plotted and each experimental run is color-coded. Statistical analysis was Kruskal-Wallis with Dunn multiple comparisons test. (D) Total fluorescence of phalloidin staining (i.e. actin filaments) on each ECM. (**** = p<0.0001, FN n=967 cells, Lam n=846 cells pooled from the same 3 independent experiments used to generate data in Figure 1B and 1C). Means and standard error of the mean for each experiment are represented with black symbols, all data points are plotted and each experimental run is color-coded. Statistical analysis was Kruskal-Wallis with Dunn multiple comparisons test. (E) Representative tracks of macrophages migrating randomly on 10 μg/mL fibronectin or laminin over a 16 hour time period, generated by the TrackMate FIJI plugin. Scale bar = 200 microns. (F) Velocity (cell speed) in microns per minute and (G) Persistence (d/T) of randomly migrating macrophages on 10 μg/mL of the indicated ECM (**** = p<0.0001, *** = p=0.0001, P-L-L n=1264 tracks, Col n=959 tracks, FN n=1004 tracks, Lam n=1559 tracks, VN n=773 tracks pooled from 2 experiments). Means and standard error of the mean for each experiment are represented with black symbols, all data points are plotted and each experimental run is colorcoded. Statistical analysis was Kruskal-Wallis with Dunn multiple comparisons test.

Given the cytoskeleton’s role in dynamic cellular processes we hypothesized further that ECM environment influences cell motility. Laminin and fibronectin were chosen as model systems to further interrogate this idea, as these ECMs influenced macrophages in such strikingly converse ways. Macrophages plated on fibronectin adopted a more mesenchymal behavior during random migration experiments, moving via coherent leading edge protrusion coupled to retraction at the cell rear (**Supplemental movie 1**). On the other hand, laminin drove macrophages to adopt a ‘chaotic’ style of migration during random migration experiments that largely dispenses with stable edge protrusion in favor of a more randomized, shorter lived protrusion cycle (**Supplemental movie 2**). As a result, migration on fibronectin gives rise to linear migration tracks, while laminin induces meandering tracks that frequently change direction (Fig. 1E). Laminin enhances macrophage migration speed, but also causes them to change direction more often (i.e. they are less directionally persistent) (Fig. 1F, 1G). Conversely, macrophages on fibronectin are relatively slow, and less likely to change direction (Fig. 1F, 1G). Though there were motility and morphology differences on collagen, vitronectin and poly-L-lysine, these all elicited mesenchymal-like motility in macrophages, similarly to FN (Fig. 1F, 1G; Supplemental Figure 1). These data together demonstrate that macrophage morphology, cytoskeletal regulation and motility are inherently tunable by the ECM microenvironment.

### Macrophage response to extracellular matrix composition is concentration and integrin-dependent

Macrophages respond in a fundamentally different fashion to fibronectin and laminin. We therefore wondered whether varying the amount or relative composition of ECM in the microenvironment would also alter macrophage cellular dynamics. Relatively low concentrations of fibronectin allow macrophages to move significantly faster and less persistently than cells at a higher concentration (Fig. 2A; **Supplemental movie 3**). Macrophages demonstrate a similar trend on the same concentration range of collagen and vitronectin (Supplemental Figure 2A, 2B). Many ECM receptors exist on the surface of macrophages, with the heterodimeric integrins being chief mediators of adhesion. Therefore, these data indicate an inverse relationship between adhesion strength and migration speed, a fundamental relationship that has been noticed before^[34, 35]^. It is also worth noting the inverse relationship between cell speed and migratory persistence in these random migration experiments, similar to Fig. 1F, 1G. Perhaps high levels of integrin engagement with FN reinforce leading edge maintenance and directional persistence at the expense of cell speed. In contrast, macrophages plated on a relatively high concentration of laminin were faster and less persistent than their counterparts plated on lower laminin concentrations (Fig. 2B; **Supplemental movie 4**). This is the opposite of what would be expected if adhesion alone determined the macrophage response to laminin. Previous studies have advanced the idea that macrophages do not adhere well to laminin despite expressing α6β1 integrin ^[36, 37]^, which canonically binds laminin. In order to test whether substrate adhesion plays a role in the laminin response, we allowed cells to spread on different concentrations of laminin and quantified how many remained after gentle washing. Macrophages on lower laminin concentrations spread relatively well (Fig. 2C), arguing that the failure to spread on 10 µg/mL laminin is a deliberate response rather than simply a failure to sense laminin. Yet there is some merit to the idea that high concentrations of laminin do affect macrophage adhesion, as more cells wash off at the high concentration than the lower ones (Fig. 2D). However, this does not fully explain the behavior of these cells. Blocking α6 integrin via neutralizing antibody lowers adhesion to 10 µg/mL laminin even further than baseline (Fig. 2E), demonstrating that integrin-based adhesion does occur in macrophages responding to laminin. Deposition of fluorescent laminin and fibronectin across these concentration ranges was noted, arguing that the differences seen here are not due to defective laminin binding to our glass surfaces (Supplemental Figure 2C). This result suggests that α6β1 may impair formation of strong focal adhesions when a sufficient threshold of laminin is present. In total, these studies indicate that macrophages use a different style of integrin-based adhesion to bind to laminin, which facilitates a different cellular response compared to RGD-based fibronectin adhesion.

**Figure 2:**
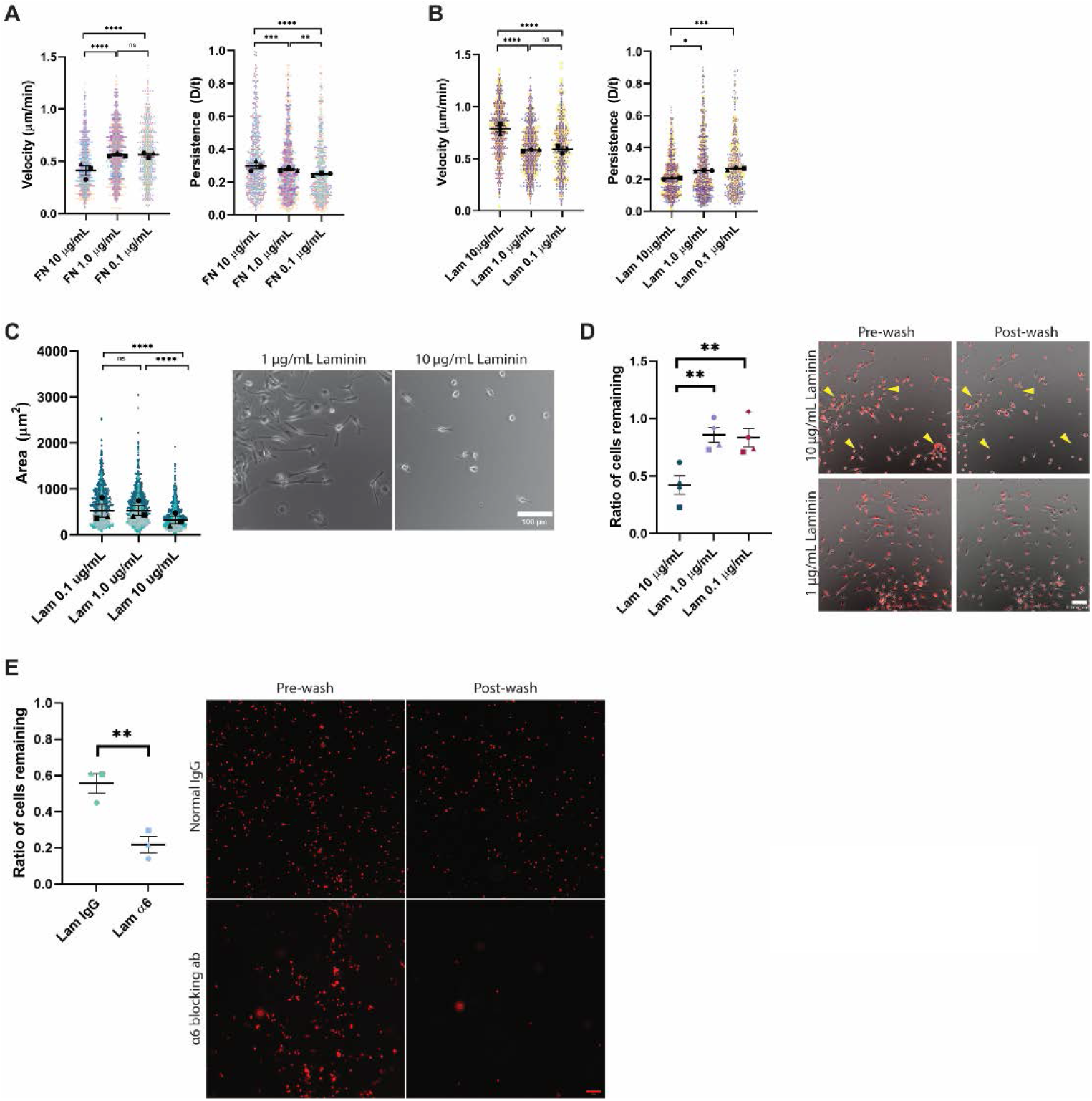
Macrophage response to extracellular matrix composition is concentration and integrin-dependent. (A) Velocity (cell speed) in microns per minute and persistence (d/T) for macrophages migrating randomly on the indicated concentration of fibronectin (FN). Means and standard error of the mean for each experiment are represented with black symbols, all data points are plotted and each experimental run is color-coded. Statistical analysis was Kruskal-Wallis with Dunn multiple comparisons test. **** = p<0.0001, *** = p=0.0004, ** = p=0.0022, ns=not significant. FN 10 μg/mL n=701 tracks, 1.0 μg/mL n=840 tracks, 0.1 μg/mL n=656 tracks. These data were pooled from three independent experiments. (B) Velocity (cell speed) in microns per minute and persistence (d/T) for macrophages migrating randomly on the indicated concentration of laminin (LAM). Means and standard error of the mean for each experiment are represented with black symbols, all data points are plotted and each experimental run is color-coded. Statistical analysis was Kruskal-Wallis with Dunn multiple comparisons test. **** = p<0.0001, *** = p=0.0002, *p=0.0182, ns=not significant. Lam 10 μg/mL n=590 tracks, 1.0 μg/mL n = 651 tracks, 0.1 μg/mL n=454 tracks. (C) Spread cell area in microns squared (*Left*), for macrophages plated on the indicated concentration of laminin (LAM). Means and standard error of the mean for each experiment are represented with black symbols, all data points are plotted and each experimental run is color-coded. Statistical analysis was Kruskal-Wallis with Dunn multiple comparisons test. **** p<0.0001, Kruskal-Wallis, Lam 0.1 μg/mL n=1439 cells, Lam 1.0 μg/mL n=1293 cells, Lam 10 μg/mL n=1279 cells pooled from 3 experiments. Representative images of 1.0 μg/mL and 10 μg/mL are shown (*Right*) for comparison. Scale bar = 100 microns. Uncropped versions of these images can be found in Supplemental Figure 4. (D) Adhesion assay for macrophages plated on the indicated concentration of laminin (LAM). Cell number was counted for each field of view before and after gentle washing. Data represents the ratio of total number of bound cells post-wash divided by the total number of cells pre-wash for each condition. Statistical analysis was done with one-way ANOVA with multiple comparisons test. Lam 10 and Lam 1.0 μg/mL **p=0.0069, Lam 10 and Lam 0.1 μg/mL **p=0.0092, N=4 independent experiments. *Right*: Representative images of tracker dye labeling superimposed on phase contrast images during indicated condition. Yellow arrowheads indicate regions where cells are present in the pre-wash condition and missing after gentle washing. Uncropped versions of these images can be found in Supplemental Figure 4. Scale bar = 100 microns. (E) Macrophages plated on 10 μg/mL Lam after 1 hour incubation at 37 °C with either IgG isotype control antibody or á6 blocking antibody. Data was analyzed and quantified identically to adhesion assay data in panel 2D. However, in this case an unpaired t-test was used to determine statistical significance, **p = 0.0084, N = 3 independent experiments. *Right*: Representative images of tracker dye labeling pre- and post-washing after exposure to IgG control or á6 blocking antibody. Scale bar = 100 microns.

### Cells change shape dramatically and ‘slingshot’ on laminin

To better understand the effects of laminin and fibronectin on macrophage cellular dynamics, we analyzed the frame to frame cell shape changes of macrophages migrating on laminin and fibronectin. Macrophages plated on laminin are highly dynamic and change morphology often, while their counterparts on fibronectin are capable of maintaining a front-rear polarity that makes it possible to migrate in straight lines for hours at a time (Fig. 3A, arrowheads). The dynamic nature of the laminin-plated cells is quantifiable, as their ability to transition rapidly between balled up (more circular) and elongated (less circular) morphologies from one frame to the next is substantially higher than macrophages plated on fibronectin (Fig. 3B). Macrophages migrating on laminin can ‘skate’ around as they randomly protrude and change direction often, as in Fig. 3A. However, these cells can also undergo a process on laminin that we refer to as ‘slingshot motility’ or ‘slingshotting’ in which macrophages elongate before releasing the cell rear and launching the cell body forward (Fig. 3C). While slingshot motility is not the only strategy macrophages use to migrate on laminin, it does appear to be uniquely prevalent on laminin and to happen rarely on fibronectin (Fig. 3D). Together, these data reveal that macrophage motility on laminin is characterized by rapid and dynamic cell shape changes that give rise to their meandering migration style, whereas macrophages on fibronectin have a stable shape that maintains directional persistence.

**Figure 3:**
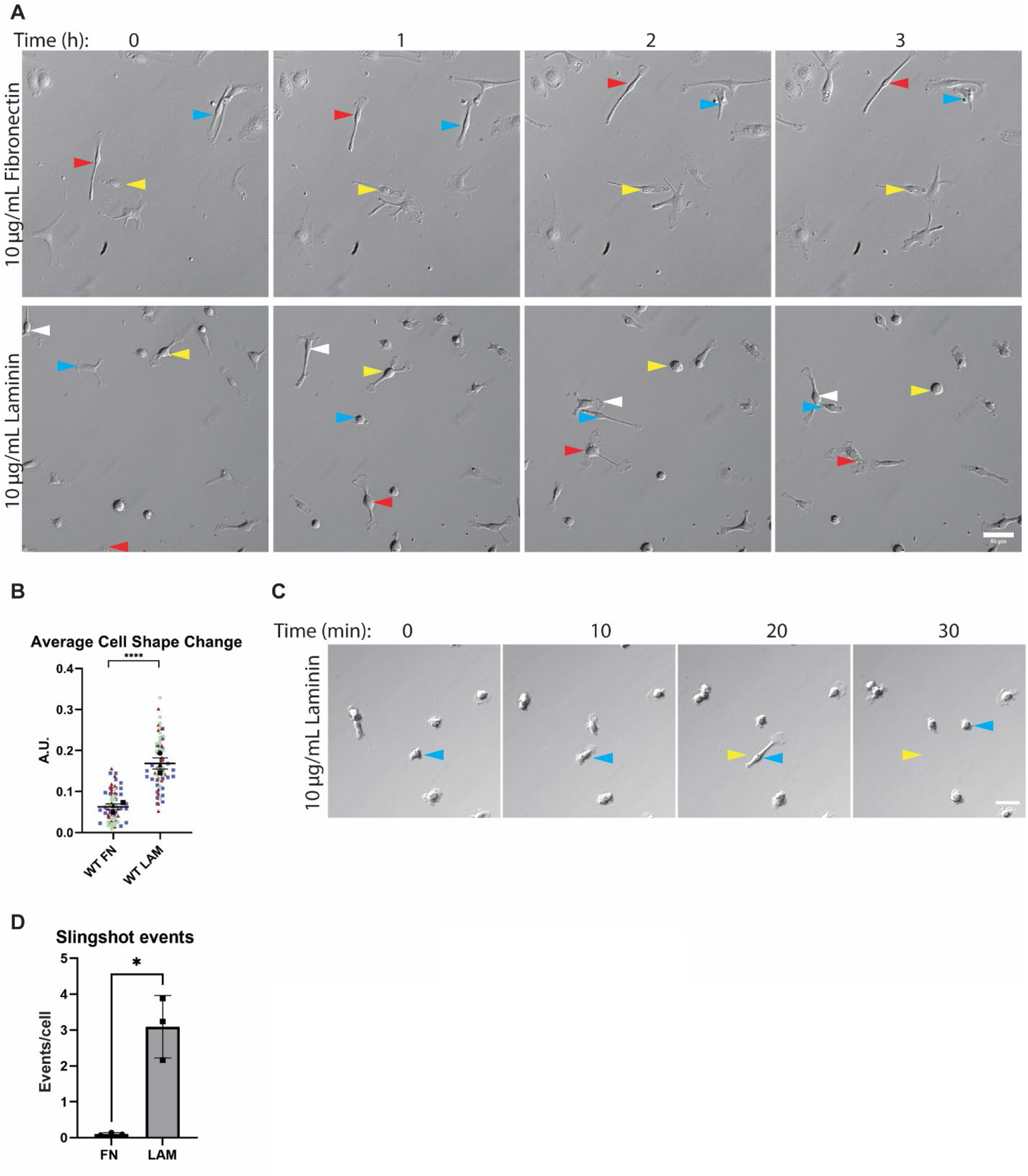
Fibronectin and laminin induce distinct modes of migration. (A) Representative relief contrast images demonstrating random migration modes of macrophages on laminin and fibronectin. Blue, red, and yellow arrowheads in relief contrast images mark different individual cells over the three hour period. This image series is part of a longer time lapse experiment. The unabridged, uncropped data can be found in Supplementary Movies 1 and 2. Images in this panel correspond to frames 1, 7, 13 and 19 of the respective movie. Scale bar = 50 microns. (B) Changes in cellular circularity occurring from frame to frame (10 minute intervals) during random motility over the course of 5 hours. Means of each experiment and standard error of the mean are represented with black symbols, all data points are plotted, and each experimental run is color-coded. ****p< 0.0001, analyzed via Mann-Whitney test. FN n = 77 cells, LAM n = 75 cells. These data were pooled from three independent experiments. (C) Representative image of a macrophage undergoing a slingshot event after plating on laminin. Blue arrowheads mark the position of the cell body in each frame. Yellow arrowheads correspond to the original position of the cell body just prior to a slingshot event. Scale bar = 40 microns. (D) Number of slingshot events per cell in each condition, when analyzed during the 5 hour intervals used to generate data in Fig. 3B. A slingshot event was defined as a change in cell shape ≥ 0.4 from one frame to the next. Events/cell was determined for each experimental run, and are plotted here with the standard error of the mean. These data were analyzed with an unpaired t-test using Welch correction. *p = 0.0269. N = three experiments.

### Non-muscle myosin II drives the rapid cell shape changes that contribute to fast motility on laminin

Several key phenotypic elements of macrophage motility on laminin resembled amoeboid motility. Their less spread appearance coupled with rapid, meandering migration suggested that these cells were not moving primarily via lamellipodial protrusions. Macrophages on laminin seemed to be using cellular contractility to skate and slingshot over the surface, rather than crawling. These observations led us to test whether myosin II contractility contributed to the laminin phenotype. Laminin-plated macrophages treated with the non-muscle myosin II inhibitor S-nitro-blebbistatin (hereafter referred to as blebbistatin) are less dynamic than control cells (Fig. 4A, **Supplementa Movie 5**). All tested concentrations of blebbistatin decreased macrophage migration speed on laminin, but had no effect on persistence (Fig. 4B, 4C). Interestingly, this corresponded with decreased dynamic cell shape changes and fewer slingshot events in the presence of blebbistatin (Fig. 4D, 4E). Macrophages plated on FN were less severely affected by blebbistatin, with only the highest blebbistatin dose slowing motility and no effect on persistence at any dose (Supplemental Figure 3A, 3B). These data indicate that myosin II activity is a fundamental contributor to the dynamic cell shape changes that drive rapid macrophage motility on laminin.

**Figure 4:**
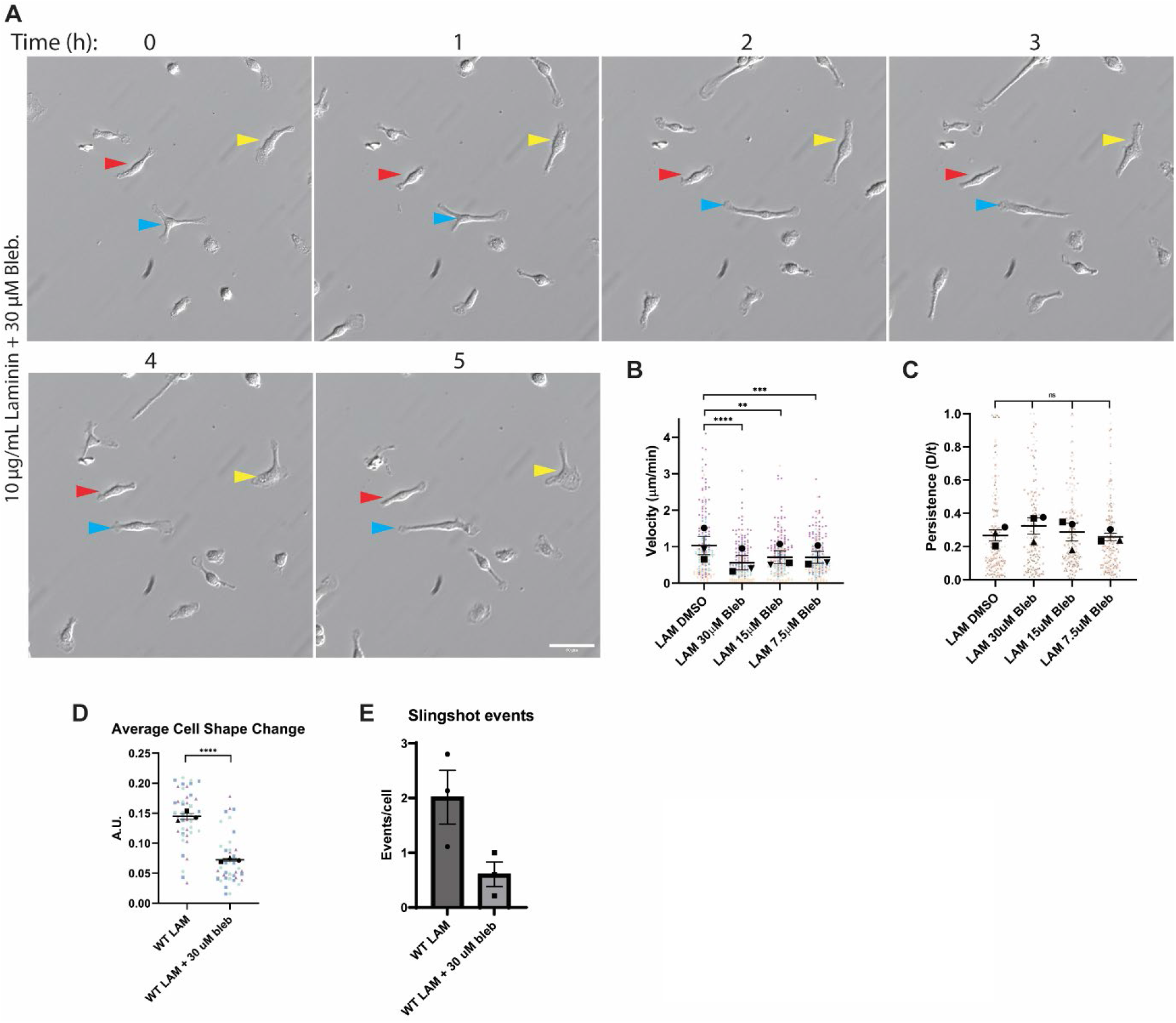
Non-muscle myosin II drives the rapid cell shape changes that contribute to fast motility on laminin. (A) Representative relief contrast images demonstrating random migration of laminin-plated macrophages migrating in the presence of 30 μM blebbistatin, which inhibits non-muscle myosin II. Blue, red, and yellow arrowheads in relief contrast images mark different individual cells over the five hour period. This image series is part of a longer time lapse experiment. The unabridged, uncropped data can be found in Supplementary Movie 5. Images in this panel correspond to frames 1, 7, 13, 19, 25 and 31 of the movie. Scale bar = 50 microns. (B) Velocity (cell speed) in microns per minute and (C) persistence (d/T) for macrophages migrating on laminin (LAM) and subjected to indicated doses of blebbistatin, or DMSO as the negative control. Means and standard error of the mean for each experiment are represented with black symbols, all data points are plotted and each experimental run is color-coded. Statistical analysis was Kruskal-Wallis with Dunn multiple comparisons test. **** = p<0.0001, *** = p=0.0004, ** = p=0.0031, ns=not significant; Lam DMSO n=200 tracks, Lam 30 μM Blebbistatin n=172 tracks, Lam 15 μM Blebbistatin n=204 tracks, Lam 7.5 μM Blebbistatin n=197 tracks pooled from 3 experiments). (D) Changes in cellular circularity occurring from frame to frame (10 minute intervals) during random motility over the course of 5 hours. Means of each experiment and standard error of the mean are represented with black symbols, and all data points are plotted and each experimental run is color-coded. ****p<0.0001, analyzed via Mann-Whitney test, Lam n=48 cells, Lam 30 μM Blebbistatin n=44 cells pooled from three independent experiments. (E) Number of slingshot events per cell in each condition, when analyzed during the 5 hour intervals used to generate data in Fig. 4D. A slingshot event was defined as a change in cell shape ≥ 0.4 from one frame to the next. Events/cell was determined for each experimental run, and are plotted here with the standard error of the mean.

### Dual inhibition of myosin II and ROCK suppresses laminin phenotype

Rho-associated protein kinase (ROCK) plays a fundamental role in modulating the actin cytoskeleton ^[38]^. Though it is a major activator of myosin contractility ^[39]^, ROCK also influences the activity of cofilin ^[38]^, Ezrin/Radixin/Moesin (ERM) proteins ^[40, 41]^, and the formin FHOD1 ^[42]^. All of these proteins have been implicated in dynamic cellular processes like motility. Since myosin inhibition alone did not raise migratory persistence on laminin, we wondered whether dual inhibition with blebbistatin and the ROCK inhibitor Y-27632 would cause macrophages on laminin to behave more like cells plated on fibronectin. Macrophage cell speed on laminin was significantly reduced by dual inhibition of myosin II and ROCK, to the extent that they were slightly slower than untreated macrophages on fibronectin (Fig. 5A, 5B; **Supplemental Movie 6**). Dual inhibition also raised migratory persistence on laminin, to the extent that this population became indistinguishable from the fibronectin control cells (Fig. 5C; **Supplemental Movie 6**). Macrophages plated on fibronectin were also slower and more persistent with dual inhibition compared to untreated cells on fibronectin (Fig. 5B, 5C; **Supplemental Movie 7**). All of these data indicate that myosin II and ROCK activity are crucial to the laminin phenotype. We hypothesized that this effect is due to laminin’s ability to activate ROCK and myosin contractility more efficiently than fibronectin. ROCK activation leads to phosphorylation of myosin regulatory light chain (MLC), leading to activation of myosin II contractility ^[43, 44]^. Additionally, ROCK activates LIM kinase ^[38]^, which in turn phosphorylates cofilin, thereby inactivating it ^[45, 46]^. Therefore, to interrogate our hypothesis further, p-MLC (Ser20) and p-cofilin (Ser3) were used as readouts for ROCK activity on fibronectin and laminin. Contrary to our expectation, phosphorylation of MLC at Ser20 and cofilin at Ser3 were not significantly different when cells were plated on the two ECMs, when normalized to GAPDH (Fig. 5D). We then wondered if the activation levels of myosin and ROCK were less important than how the two proteins localized within macrophages when interacting with each ECM component. GFP-myosin IIA knock-in macrophages were subjected to Airyscan imaging. 3D reconstructions revealed a wider band of GFP-myosin IIA throughout laminin-plated macrophages, whereas cells on fibronectin tended to localize GFP-myosin IIA to a more restricted area at the bottom of the cell (Fig. 5E, white brackets). We sought further quantitative evidence for this localization and used myosin IIA and p-MLC staining in fixed cells imaged with standard confocal microscopy. Indeed, we demonstrate that a greater fraction of the total myosin IIA and p-MLC signal is present at the bottom (ventral surface) of FN-plated cells (Fig. 5F), reflecting an alteration in active myosin localization toward the adhesive plane in these cells. Though myosin II and ROCK clearly contribute to the laminin phenotype, we find no evidence for hyperactivation of contractile signaling. Instead, the contractile apparatus that is normally in close proximity to the integrin-containing bottom of the cell (on FN) is redistributed on laminin and appears to be spread throughout the cell cortex rather than confined at the bottom of the cell.

**Figure 5:**
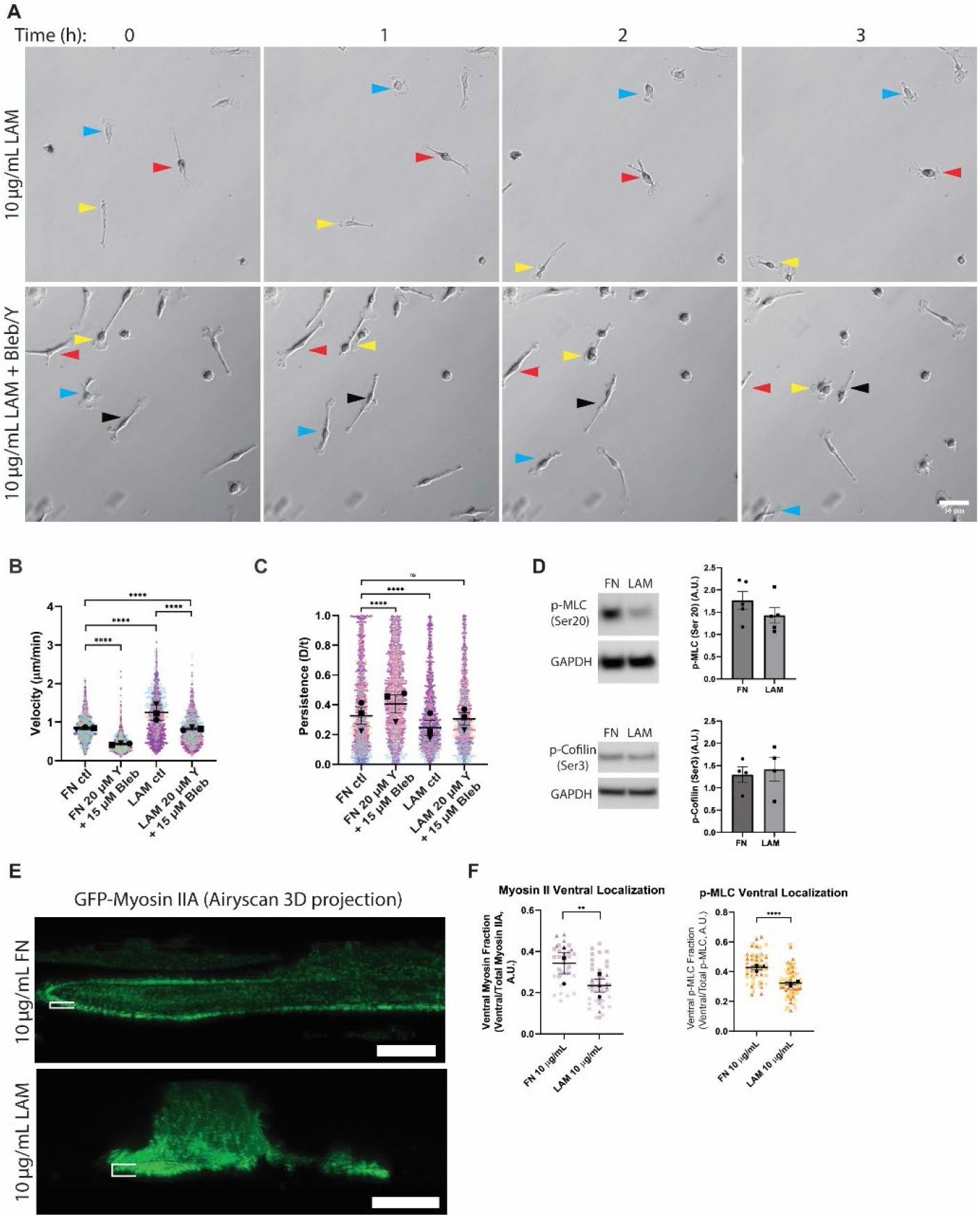
Dual inhibition of myosin II and ROCK suppresses laminin phenotype. (A) Representative relief contrast images demonstrating random migration of laminin-plated macrophages treated with DMSO or a combination of 20 μM Y-27632 (ROCK inhibitor, ‘Y’) and 15 μM Blebbistatin (‘Bleb’). Blue, red, and yellow arrowheads in relief contrast images mark different individual cells over the three hour period. These are part of a longer time lapse experiment. The unabridged, uncropped images of the dual inhibitor condition can be found in Supplementary Movie 6. Images in this figure correspond to frames 25, 31, 37 and 43 of Supplemental Movie 6. Scale bar = 50 microns. (B) Velocity (cell speed) in microns per minute and (C) persistence (d/T) for macrophages migrating on fibronectin (FN) or laminin (LAM) and subjected to DMSO (ctl) or 20 μM Y-27632 (‘Y’) + 15 μM Blebbistatin (‘Bleb’). Means and standard error of the mean for each experiment are represented with black symbols, all data points are plotted and each experimental run is color-coded. Statistical analysis was Kruskal-Wallis with Dunn multiple comparisons test. **** = p<0.0001, *** = p=0.0005, ns=not significant; FN ctrl n=1826 tracks, FN 20 μM Y-27632 15 μM Bleb n=1779 tracks, Lam ctrl n =1712 tracks, Lam FN 20 μM Y-27632 15 μM Bleb n=1539 tracks pooled from three experiments. (D) *Left*: Western blot images of p-myosin light chain (p-MLC, Ser20), and p-cofilin (Ser3). GAPDH is included as a loading control. *Right*: Quantification of p-MLC and p-cofilin signal after normalization to GAPDH. N = 5 for p-MLC and N = 4 for p-cofilin. Bars represent standard error of the mean. Uncropped images of these blots can be found in Supplemental Figure 5. (E) 3D Airyscan projections of GFP-myosin IIA knock-in macrophages plated on 10 μg/mL FN or LAM. White brackets denote z-width of GFP-myosin IIA band in each condition. Scale bar = 10 microns. (F) Quantification of myosin IIA (*Left*) and p-MLC (*Right*) fluorescence level in the lowest confocal z-slice relative to the total amount of fluorescence staining in the cell. Means and standard error of the mean for each experiment are represented with black symbols, all data points are plotted and each experimental run is color-coded. Statistical analysis was evaluated with an unpaired t-test. **p-value = 0.0054, ****p-value < 0.0001. n = 37 cells for myosin II on FN, 49 cells for myosin II on LAM, 66 cells for p-MLC on FN, and 78 cells for p-MLC on LAM, pooled from three experiments.

### Macrophages preferentially sense laminin in mixed ECM conditions

Macrophages respond to a laminin-rich environment by utilizing myosin contractility to establish a rapid, meandering mode of migration. We next sought to determine whether this response was a passive one arising from an inability to adhere well to laminin, or a deliberate response to a laminin-rich environment. Fibronectin induces strong adhesion and spreading in macrophages, so its inclusion alongside laminin should suppress the laminin phenotype if the latter arises due to impaired adhesion. In contrast to this expectation, we saw that the migratory characteristics of macrophages in the mixed ECM context was largely influenced by how much laminin was present. Macrophages migrate faster and less persistently in a laminin dose-dependent fashion when fibronectin is held at our baseline concentration of 10 µg/mL (Fig. 6A; **Supplemental Movies 8-9**). On the other hand, when laminin is held constant at 10 µg/mL, both migration speed and persistence are largely resistant to increasing concentrations of fibronectin (Fig. 6B). At the 10 ug/mL LAM + 30 ug/mL FN macrophages begin to slow slightly in response to fibronectin, but their motility is still higher than baseline FN migration (Fig. 6B; **Supplemental Movie 10**). One interpretation of this data would be that macrophages are physically prevented from interacting with fibronectin when laminin is present. When fluorescently labeled ECMs (Rhodamine-FN and HiLyte 488-LAM) are used in this assay (both at 10 µg/mL) we are able to detect a strong intracellular Rhodamine-fibronectin localization in the presence of laminin (Fig. 6C), suggesting that macrophages in this context can still interact with fibronectin. Both ECM components were functionalized well on glass when mixed (Supplemental Figure 3C), arguing against the interpretation that the mixed ECM phenotype is due to only laminin being effectively deposited on the surface. These findings support the notion that interactions between specific integrins and distinct elements of the ECM can be sufficient to drive motile behavior by differentially regulating cytoskeletal organization and function.

**Figure 6:**
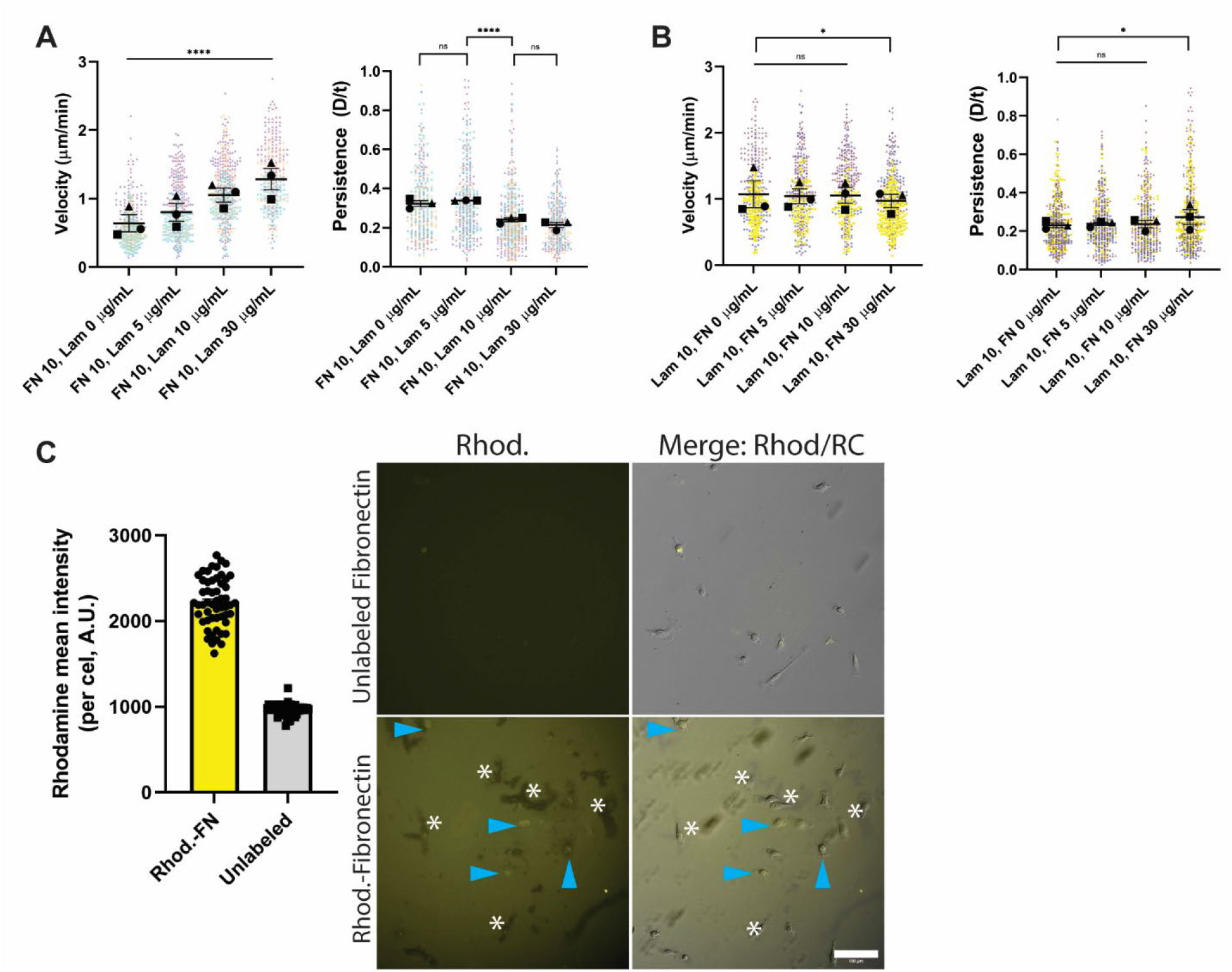
Macrophages preferentially sense laminin in mixed ECM conditions. (A) Velocity (cell speed) in microns per minute and persistence (d/T) for macrophages migrating on fibronectin or fibronectin + laminin at various concentrations. Means and standard error of the mean for each experiment are represented with black symbols, all data points are plotted and each experimental run is color-coded. Statistical analysis was Kruskal-Wallis with Dunn multiple comparisons test. ****p<0.0001, ns=not significant; FN 10 Lam 0 μg/mL n=360 tracks, FN 10 Lam 5 μg/mL n=374 tracks, FN 10 Lam 10 μg/mL n=460 tracks, FN 10 Lam 30 μg/mL n=335 tracks, pooled from 3 experiments. (B) Velocity (cell speed) in microns per minute and persistence (d/T) for macrophages migrating on laminin or laminin + fibronectin at various concentrations. Means and standard error of the mean for each experiment are represented with black symbols, all data points are plotted and each experimental run is color-coded. Statistical analysis was Kruskal-Wallis with Dunn multiple comparisons test. *p=0.0124, for persistence, *p = 0.0273, for velocity; Lam 10 FN 0 μg/mL n=424 tracks, Lam 10 FN 5 μg/mL n=393 tracks, Lam 10 FN 10 μg/mL n=394 tracks, Lam 10 FN 30 μg/mL n=456 tracks, pooled from 3 experiments. Examples of cells migrating on mixed ECMs can be found in Supplemental Movies 8-10. (C) *Left*: Rhodamine fluorescence intensity of macrophages plated on Rhodamine-fibronectin or unlabeled fibronectin. *Right*: Representative images of Rhodamine intensity, alongside Rhodamine/relief contrast merge. Blue arrowheads denote macrophages that are Rhodamine+, while white asterisks highlight areas where fibronectin appears to have been ripped away. Scale bar = 100 microns. Uncropped versions of these images can be found in Supplemental Figure 4.

### Modulation of RGD-based adhesion is sufficient to shift cells on FN to LAM-like phenotype

The results from mixed ECM experiments indicate that modulating integrin engagement alone may be sufficient to shift cellular behavior through differential cytoskeletal regulation. We decided to test this idea further by determining whether disruption of RGD-based adhesion to fibronectin rendered cells in that context more ‘laminin-like’. The small molecule cilengitide was utilized in these experiments due to its ability to disrupt a broad range of RGD-binding integrins, most notably αv-containing integrin pairs ^[47]^. Time lapse experiments demonstrated that cilengitide-treated macrophages were small and highly dynamic compared to their untreated counterparts on FN (Fig. 7A; **Supplemental Movie 11**). These experiments revealed a significant dose-dependent increase in cell speed and decrease in persistence in cilengitide-treated cells compared to control macrophages on FN (Fig. 7B, 7C), reminiscent of the laminin motility phenotype. Furthermore, cilengitide-treated macrophages on FN showed a laminin-like decrease in spread cell area and shift toward a more circular morphology (Fig. 7D, 7E). Frame to frame macrophage cell shape changes on FN were increased by cilengitide treatment, though in a much more modest fashion than elicited by laminin (Fig. 7F), and cilengitide treatment did not cause macrophages to initiate slingshot motility on fibronectin (Fig. 7G). Despite striking similarities, these data demonstrate that cilengitide treatment on fibronectin does not perfectly phenocopy laminin. In addition, the effect of cilengitide cannot be chalked up to altered adhesive strength, as cilengitide does not impair macrophage adhesion to fibronectin (Fig. 7H). This suggests that there are other FN-binding integrins or integrin-independent adhesion systems that were not targeted by cilengitide and can mediate adhesion without stimulating spreading. Taken together, our data suggest that it is possible for macrophages to tune their migratory behavior according to which surface receptors are interacting with the ECM microenvironment.

**Figure 7:**
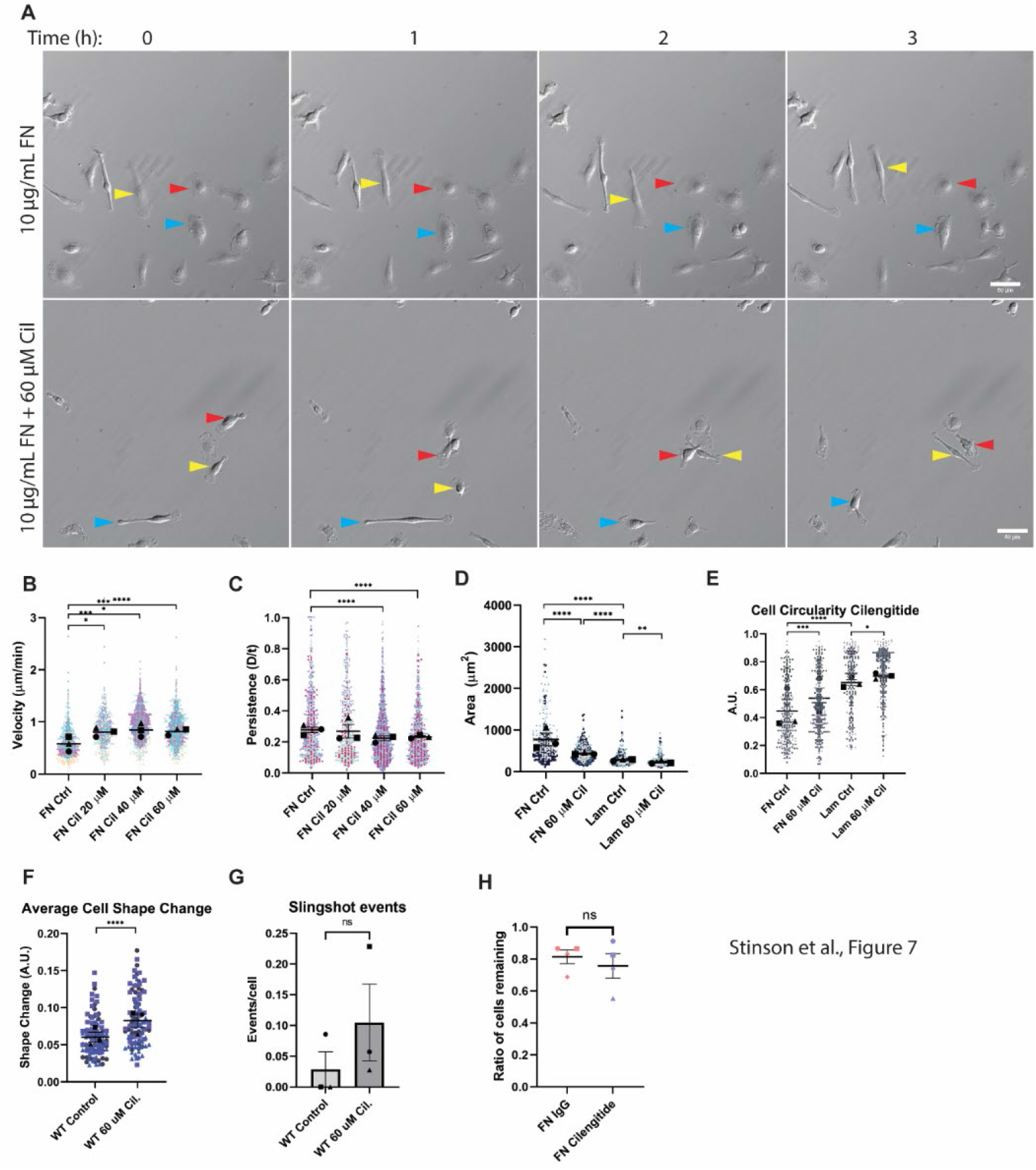
Modulation of RGD-based adhesion is sufficient to shift cells on FN to LAM-like phenotype. (A) Representative relief contrast images demonstrating random migration of fibronectin-plated macrophages treated with DMSO or 60 μM Cilengitide (‘Cil’), an RGD-binding integrin inhibitor. Blue, red, and yellow arrowheads in relief contrast images mark different individual cells over the three hour period. These are part of a longer 16 hour time lapse experiment. The unabridged, uncropped data can be found in Supplementary Movie 11. Images in this panel correspond to frames 33, 39, 45 and 51 of the movie. Scale bar = 50 microns. (B) Velocity (cell speed) and (C) persistence (d/T) for macrophages migrating on fibronectin (FN) in the presence of the indicated dose of Cilengitide. Means and standard error of the mean for each experiment are represented with black symbols. All data points are plotted and each experimental run is color-coded. Statistical analysis was Kruskal-Wallis with Dunn multiple comparisons test. ****p<0.0001; FN Ctrl n=716 tracks, FN Cil 20 μM n=463 tracks, FN Cil 40 μM n=1696 tracks, FN Cil 60 μM n=883 tracks, pooled from 3 experiments. (D) Spread cell area of macrophages on FN or LAM ± 60 μM Cilengitide. Means and standard error of the mean for each experiment are represented with black symbols. All data points are plotted and each experimental run is color-coded. Statistical analysis was Kruskal-Wallis with Dunn multiple comparisons test. **** = p<0.0001, ** = p=0.0060; FN Ctrl n=315 cells, FN Cil 60 μM n=359 cells, Lam Ctrl n=255 cells, Lam Cil 60 μM n= 277 cells pooled from 3 experiments. (E) Circularity values for macrophages on fibronectin or laminin ± 60 μM Cilengitide. Means and standard error of the mean for each experiment are represented with black symbols. All data points are plotted and each experimental run is color-coded. Statistical analysis was Kruskal-Wallis with Dunn multiple comparisons test. **** = p<0.0001, *** = p=0.0002, * = p=0.0406; FN Ctrl n=315 cells, FN Cil 60 μM n=359 cells, Lam Ctrl n=255 cells, Lam Cil 60 μM n=277 cells, pooled from 3 experiments. (F) Changes in cellular circularity occurring from frame to frame (10 minute intervals) during random motility over the course of 5 hours. Means of each experiment and standard error of the mean are represented with black symbols, and all data points are plotted and each experimental run is color-coded. Statistical analysis was Mann-Whitney. ****p<0.0001; n = 106 cells pooled from 3 experiments. (G) Number of slingshot events per cell in each condition, when analyzed during the 5 hour intervals used to generate data in Fig. 7F. Events/cell was determined for each experimental run, and are plotted here with the standard error of the mean. (H) Macrophages plated on 10 μg/mL FN were subjected to gentle washing after treatment with IgG or 60 μM Cilengitide. Data is reported as ratio of cells remaining after wash (1 = 100% adherence). An unpaired t-test was used to determine statistical significance. p-value = n.s., not significant, N = 4 independent experiments.

Myosin II and ROCK are major contributors to the high motility/low persistence laminin phenotype. We next asked whether cilengitide’s ability to increase motility and suppress persistence on fibronectin was also due to these factors. ROCK inhibition corrected the FN cilengitide phenotype most dramatically. Treatment with 20 µM Y compound suppressed motility in the presence of cilengitide and returned persistence values back to WT levels (Fig. 8A-8C; **Supplemental Movie 12**). ROCK inhibition with Y compound also allowed cilengitide-treated cells to spread and become less circular (Fig. 8D, 8E). As with similar phenotypes on laminin, the cilengitide migration and morphology phenotypes on fibronectin appear to require ROCK activity. Treatment with 15 µM blebbistatin, a dose that does not impact macrophage motility on fibronectin at baseline (Supplemental Fig. 3), also impacted the cilengitide phenotype. As we anticipated, treatment with blebbistatin in the presence of cilengitide brought macrophage cell speed down to control FN levels, but had little effect on persistence (Fig. 8A-8C; **Supplemental Movie 13**). In addition, blebbistatin did not spread or elongate cilengitide-treated macrophages on FN (Fig. 8D, 8E). These outcomes are similar to the effect of blebbistatin on laminin migration. We conclude based on these results that impairing RGD-integrin interactions causes macrophages on FN to become more laminin-like, adopting morphology and motility phenotypes that require myosin II and ROCK activity.

**Figure 8:**
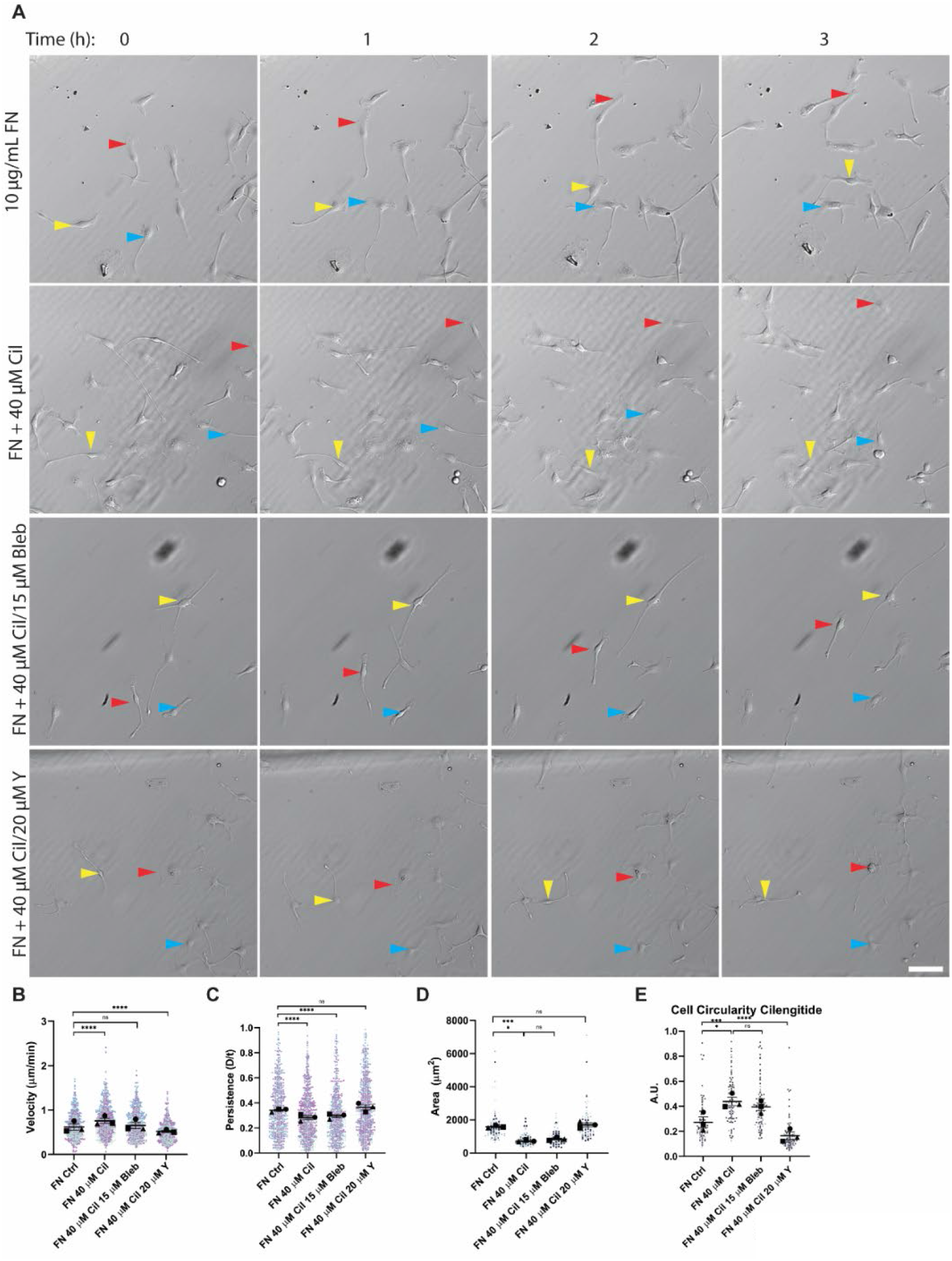
Inhibition of myosin II and ROCK ameliorates cilengitide’s effect on FN motility. (A) Representative relief contrast images demonstrating random migration of fibronectin-plated macrophages treated with DMSO, 40 μM Cilengitide (‘Cil’), 40 μM Cilengitide (‘Cil’) + 15 μM Blebbistatin, or 40 μM Cilengitide (‘Cil’) + 20 μM Y-27632 (‘Y’). Blue, red, and yellow arrowheads in relief contrast images mark different individual cells over the three hour period. These are part of a longer time lapse experiment. The unabridged, uncropped data can be found in Supplemental Movies 12 and 13. Images in this panel correspond to frames 1, 7, 13 and 19 of the movies. Scale bar = 100 microns. (B) Velocity (cell speed) and (C) persistence (d/T) for macrophages migrating on fibronectin (FN) in the presence of the indicated inhibitor. Means and standard error of the mean for each experiment are represented with black symbols. All data points are plotted and each experimental run is colorcoded. Statistical analysis was Kruskal-Wallis with Dunn multiple comparisons test. **** = p<0.0001, FN Ctrl n= 735 tracks, FN Cil n=891 tracks, FN Cil + Bleb n=696 tracks, FN Cil + Y n=798 tracks, pooled from 3 experiments. (D) Spread cell area of macrophages on FN, in presence of indicated inhibitor. Means and standard error of the mean for each experiment are represented with black symbols. All data points are plotted and each experimental run is color-coded. Statistical analysis was Kruskal-Wallis with Dunn multiple comparisons test. **** = p<0.0001, FN Ctrl n= 105 cells, FN Cil n=105 cells, FN Cil + Bleb n=105 cells, FN Cil + Y n=103 cells, pooled from 3 experiments. (E) Circularity values for macrophages on fibronectin in presence of indicated inhibitor. Means and standard error of the mean for each experiment are represented with black symbols. All data points are plotted and each experimental run is color-coded. Statistical analysis was Kruskal-Wallis with Dunn multiple comparisons test. **** = p<0.0001, FN Ctrl n= 105 cells, FN Cil n=105 cells, FN Cil + Bleb n=105 cells, FN Cil + Y n=103 cells, pooled from 3 experiments.

## DISCUSSION

Macrophages are known to express a variety of integrins that bind different ECM components *in vivo*, though the influence of each ECM component on macrophage signaling is not fully known. There are many competing signals in a local tissue microenvironment that influence the wide range of macrophage behaviors, especially in the context of trauma and infection.^[48, 49]^ Soluble chemotactic cues and substrate stiffness are well established as factors that influence macrophage migration.^[50, 51]^ However, dense tissue environments *in vivo* house a complex array of signaling cues, including shifts in ECM composition. The present study demonstrates that macrophages adopt distinct shapes and sizes that alter their migratory behavior based solely on ECM composition. If a specific concentration of fibronectin or laminin is sufficient to alter the motility and morphology of macrophages *in vitro*, then fibrous ECM glycoproteins *in vivo* may synergize with or impair other migratory cues, such as cytokines and chemokines present in the same microenvironment. The large array of integrins expressed by macrophages may give rise to differential signaling based on how a macrophage ‘reads’ the ECM in its microenvironment. The present study offers the distinction between how macrophages respond to fibronectin and laminin as a proof of principle that this process does occur and is sufficient to differentially regulate the actomyosin cytoskeleton, leading to ECM-specific alterations in cellular structure and dynamics.

Macrophages responding to a concentration gradient of laminin or fibronectin beautifully demonstrate the ability of the cell to sense its ECM environment. Fibronectin induces strong adhesion in many cell types.^[52, 53]^ Therefore, high concentrations make cells so adherent that their cell speed decreases, while integrins are less engaged at lower concentrations of fibronectin, resulting in lower adhesion and freer movement. This relative effect on cell speed is observed in the present study. Additionally, we reveal that macrophages tend to migrate more persistently as cell speed decreases. To our surprise, laminin seems to elicit a different trend in macrophages. We observed that higher concentrations of laminin cause faster, less directionally persistent motility in macrophages while lower concentrations stimulate slower, more persistent motility. Our adhesion experiments argue that α6β1 integrin is the primary receptor for laminin binding and adhesion in macrophages. Macrophage expression of α6β1 integrin is also supported by other studies ^[18, 54]^. Macrophages express numerous fibronectin-binding integrins, so one interpretation of these data is that macrophages activate less integrin signaling when plated on laminin compared to fibronectin, which could lead to an inherent difference in adhesion. This could explain why macrophages are faster and less adherent at relatively high concentrations of laminin compared to fibronectin. A previous study using neuronal cells suggested that laminin might be able to impair focal adhesion formation ^[55]^. This is a compelling idea worthy of further study, but it does not explain why macrophages adhere better, spread more and migrate more slowly on lower concentrations of laminin compared to higher ones. We suspect that the macrophage response to high laminin levels is a deliberate choice, and that specific signaling events are stimulated by α6β1-laminin ligation. There may be a threshold that must be reached in order for this high motility/low persistence migratory style to be enacted. It is also possible that α6β1 integrin-independent adhesion also influences the cellular response to high or low levels of laminin.

Macrophages plated on laminin migrate differently than on fibronectin. We sought a quantitative framework for understanding how single cell dynamics over the course of minutes are altered within our overnight time lapse experiments. Our quantitative strategy could also be used to ascertain whether specific perturbations to the actomyosin cytoskeleton alter dynamic cell shape changes, and whether such changes contribute to the differential response on laminin and fibronectin. The cellular response to laminin is strikingly different than the response to any other ECM we tested. At any single time point, macrophages migrating on laminin will be poorly spread. They are capable of retaining this morphology and migrating randomly in a behavior best described as ‘hovering’ or ‘skating’. However, laminin also induces acute spreading events that generally last several frames (i.e. 10 to 30 minutes), followed by a translocation capable of propelling the cell relatively long distances. When the circularity of macrophages on laminin is measured frame-by-frame, there is a significant change indicating a rapid transition between balled-up and elongated morphology. We describe this behavior as ‘slingshot motility’ because it suggests a process wherein gathering tension is suddenly released to allow an object to shoot forward at a rapid pace. Macrophages do not maintain directional polarity after ‘firing’ themselves forward. They either adopt a localized exploration phase in which they skate randomly, or take off immediately in a random direction via another slingshot event. The contractile appearance of the laminin-plated macrophages, and the slingshot motility that looks reminiscent of myosin II-dependent mesenchymal cell retraction, led us to hypothesize that contractility plays an important role in the laminin response. Indeed, myosin II is required for the rapid cell shape change and slingshot behavior seen on laminin. However, blebbistatin treatment did not cause macrophages to spread on laminin. Fast migration on laminin was dependent upon myosin II, but cells retained their relatively low persistence in the presence of blebbistatin. Instead, ROCK inhibition was required on top of myosin II inhibition to shift cells on laminin toward a more ‘FN-like’ phenotype. These findings suggest that there may be myosin II-independent features of ROCK’s contribution to the laminin phenotype. ROCK is known to activate specific formins^[42]^, LIM kinase (which inhibits cofilin)[38, 45, 46], and ERM-domain containing proteins^[40, 41]^. Of particular interest, the ERM protein ezrin links actin to the plasma membrane and must be removed before Arp2/3 complex can be recruited to form a lamellipodial protrusion ^[56]^. ROCK could promote linear actin filaments, downregulate their disassembly, and further stabilize them at the plasma membrane distinct from ROCK’s role in activating myosin II-dependent contractility. Perhaps inhibiting ROCK counteracts these processes and allows cells to spread, and the morphological change is sufficient to raise migratory persistence.

ROCK and myosin II were strongly implicated in the high speed/low persistence laminin migratory phenotype. It is straightforward to hypothesize that α6β1 integrin-laminin ligation strongly stimulates ROCK/myosin II compared to fibronectin, where adhesion does not require α6β1 integrin, macrophages readily spread, and motility is not so contractility-dependent. We used both Ser20 phosphorylation of myosin regulatory light chain and Ser3 phosphorylation of cofilin as readouts for myosin contractility and ROCK activity. However, to our surprise there is no discernible difference in phosphorylation on LAM compared to FN. This finding was highly counterintuitive at first. However, the combination of weaker adhesion (via α6β1 integrin) and altered actomyosin structures could be sufficient to shift motility on LAM without greatly enhancing the overall contractile signaling downstream of receptor engagement (i.e. MLC phosphorylation). Indeed, we show support for the first part of this statement via adhesion assays showing that while α6β1 integrin is required for laminin-based adhesion, the overall strength of this interaction is weaker than RGD-binding integrins interacting with fibronectin. We also find support for the notion that actomyosin structures are organized differently when macrophages are plated on LAM versus FN. Macrophages on fibronectin are already known to be strongly adherent. Myosin II plays a major role in forming focal adhesions and generating traction for cells on fibronectin.^[57]^ In line with this work, myosin and p-MLC are both more confined toward the bottom of macrophages plated on FN while macrophages on LAM have a significantly wider distribution of these proteins in the z-dimension. This suggests that ECM composition may control the cellular localization of ROCK and myosin more than their activity by sequestering myosin near the cortex when laminin is bound by α6β1, rather than at the adhesive plane of the cell. Future measurements of cortical and membrane tension in FN- and LAM-plated macrophages might be insightful, as cortical contractility and tension may play a major role in the LAM motility phenotype.

Since laminin- and fibronectin-based adhesion require distinct integrin pairs, we set out to determine whether one or the other could exert dominance in an experiment where we made both ECM proteins available for adhesion. We expected that the relatively strong adhesion imparted by RGD-binding integrins interacting with fibronectin would supersede the relatively weak adhesion imparted by α6β1. To our surprise, macrophages plated in these conditions seem to ignore fibronectin and respond exclusively to the laminin present, as judged by their motility and morphology. This response is surprisingly durable, as we have not been able to demonstrate a concentration of fibronectin that ‘wins’ over laminin. Furthermore, the response increases linearly as a function of the laminin concentration present in the mixture. Fluorescent fibronectin and laminin co-assembled on the same glass surface were both readily detectable simultaneously, so the result cannot be chalked up to one ECM being more efficiently deposited on the glass than the other. Our evidence also suggests that laminin isn’t physically hindering fibronectin from being detected. Macrophages can tear the fibronectin away from the surface and internalize it, potentially through a Caveolin-1-dependent process.^[58, 59]^. These data lead us to propose a model wherein laminin retains its ‘dominance’ in this context via inhibitory crosstalk with RGD-binding integrins. Previous efforts in neuronal cells have demonstrated a similar dominance of laminin over other ECMs, which the authors suggest could be due to inhibition of focal adhesions ^[55]^. This could be enforced at the level of signaling factors downstream of integrin that control myosin II and ROCK activity and localization. The full signaling capacity of the many isoforms of laminin is not fully known. Laminin 511 and 411, for example, have been shown *in vivo* to alter monocyte differentiation as monocytes extravasate and pass through the laminin-rich endothelial monolayer and basement membrane.^[60]^ Whatever the case, our study presents further evidence that the laminin phenotype arises from an active process initiated by α6β1-LAM ligation rather than a passive process related to less robust adhesion.

We turned our attention to fibronectin motility after having determined that myosin II and ROCK are important for the macrophage response to laminin, and laminin’s dominance over fibronectin. We found that blocking RGD-binding integrins with cilengitide drove FN-plated macrophages to adopt a morphology and motility similar to cells plated on laminin, though it did not perfectly recapitulate the LAM phenotype in all respects. However, the data does point toward the idea that cells exist on a continuum and can rapidly adjust their behavior. Cilengitide shifts macrophages toward the ‘laminin side’ of the continuum. Whether due to integrin crosstalk or small molecule inhibition, these findings reinforce the notion that modulation of integrin-based adhesion is a fundamental strategy for shifting cellular behavior in complex ECM microenvironments. One molecular explanation for the cilengitide result could be that the RGD-binding integrins normally recruit ROCK and myosin II to the cell surface instead of the cortex. When inhibited with cilengitide, cortical localization of ROCK and myosin II could increase, leading to the fast, non-persistent migration and impaired cell spreading seen in this condition. We next wanted to test the idea that ROCK and myosin II are important for the ‘laminin-like’ phenotype of cilengitide-treated macrophages plated on fibronectin. Treatment of cilengitide-treated, FN-plated macrophages with blebbistatin or Y-27632 caused cells to revert back toward looking more ‘FN-like’ again. Interestingly, myosin II inhibition alongside cilengitide treatment only affected cell speed, leaving migratory persistence, cell shape and cell size unaffected. This is notable because blebbistatin had a nearly identical phenotypic effect on laminin-plated cells. ROCK inhibition had a much more general impact on the cilengitide-treated cells, reversing every phenotype induced by cilengitide. These data demonstrate that differential integrin binding or inhibition can shift cells along a continuum of phenotypic migratory and morphological responses. One major downstream effector that determines a cell’s place on this continuum is how it regulates myosin II and ROCK activity and/or localization.

One major future direction for this work will be its application to *in vivo* physiological and pathological processes. It is difficult, if not impossible, to accurately measure the exact local concentration of an individual ECM component in a tissue *in vivo*. However, ECM composition *in vivo* does have some well understood trends that we can use to our advantage. For example, the basement membrane of epithelial cells contains high concentrations of laminin, whereas the connective tissue interstitium contains fibronectin. It may be possible to track cellular responses *in vivo* or *ex vivo* and correlate them with proximity to spaces known to have a particular ECM context. These studies could reveal that there are subpopulations of macrophages that look similar from a transcriptional perspective, but respond very differently to external stimuli due to their being surrounded by different ECM contexts. Though part of the same lineage, monocytes and macrophages have distinct integrin profiles that can serve as differentiation markers, though macrophages and monocytes both express laminin-binding integrins, so it is possible that both cell types could respond in a similar fashion to FN and LAM.^[61]^ Monocytes extravasating to a site of infection or trauma will be confronted with a laminin-rich basement membrane surrounding the vasculature and separating it from the underlying connective tissue. The laminin-rich environment could drive monocytes to become more compact, adhere less tightly, and squeeze through the basement membrane to invade the interstitial space. Once in the fibronectin-rich interstitial space containing fibronectin, collagen and other fibrous ECMs, they use the FN-rich microenvironment to move in a directionally persistent fashion to their sites of action. It seems fitting that the inherently adaptable macrophage would be capable of immediately adapting to new ECM environments, presumably without requiring transcriptional changes. It remains to be seen how ECM sensing alters other elements of macrophage function beyond motility, and whether evidence can be found *in vivo* of cells undergoing ECM-sensitive behavior shifts.

Taken as a whole, the results in the present study suggest that macrophages are capable of detecting the amount and type of ECM present in their microenvironment, which leads to differential cellular responses governed by altered adhesion and cytoskeletal regulation. These cellular characteristics are controlled by differential integrin binding and downstream signaling to Myosin II and ROCK. Yet, several questions remain. What controls actomyosin organization and ROCK signaling downstream of integrins? Rho, Rac1, and CDC42 are good candidates, as all of these Rho family GTPases function as molecular switches that alter actomyosin structure and dynamics, tune migration, and alter macrophage shape.^[62, 63]^ How does ECM composition alter inflammatory responses? It stands to reason that differential integrin engagement and downstream cytoskeletal organization could allow macrophages to tune their inflammatory capacity to local ECM microenvironments. Fibronectin is well characterized as an ECM heavily associated with wound healing and repair.^[64, 65]^ However, the literature suggests that the FN isoform matters, as some are associated with inflammation while others have an anti-inflammatory influence [23, 24, 65]. Recent publications have also demonstrated that macrophages are primed to become inflammatory when interacting with fibrinogen, but when fibrinogen is cleaved to form fibrin they are primed to be anti-inflammatory ^[17]^. There is clear precedence for ECM-dependent immunomodulation, though there is likely much mechanistic detail yet to be uncovered. As the cytoskeleton is intimately involved in sensing ECM, it is highly likely that these immunomodulatory shifts can be linked to differential cytoskeletal regulation. The combination of ECM fibers, chemokines, cytokines, and other physiological immunomodulatory cues could potentially compete or enhance one another’s activity in any *in vivo* context. The relationship between macrophages and tumor microenvironments is another place where these ECM-dependent shifts could be important. The commercially available laminin 111 used in the current project (and by many other researchers) is purified from tumor cells *in vitro*. It is reasonable to speculate that aberrant ECM expression in this context may be part of what shifts the tumor immune environment toward a tumor-permissive status, and may even contribute to tumor invasion via lowered adhesion and heightened contractility.

## MATERIALS AND METHODS

### Bone marrow-derived macrophages

Hematopoeitic cells were isolated from mouse femurs. Bone marrow was flushed from the bone, filtered to prepare single cell suspensions, and then cultured in ‘macrophage media’ composed of 70% DMEM complete (containing a final concentration of 10% Fetal Bovine Serum, 1% penicillin/streptomycin, and 1% Glutamax) + 30% L929-conditioned media (for a source of M-CSF) for at least 7 days prior to experiments at 37°C, 90% humidity, and 5% CO_2_. Differentiated bone marrow macrophages were sorted from this initial bulk population on the basis of F4/80 positivity using an Alexa 488-conjugated antibody and FACS sorting. For all experiments contained in the present study, macrophages were maintained in macrophage media at 37°C, 90% humidity, and 5% CO_2_. Macrophages were passaged at 60-90% confluency. They were washed once with 1X PBS, then incubated at 4°C for 10 minutes with pre-chilled 0.5 mM EDTA. At the end of the incubation EDTA was removed and macrophages were scraped directly into 1-3 mL macrophage media (according to desired confluency after passage) and re-plated for passaging and/or experimentation.

### Mouse lines used to derive macrophages

The bone marrow-derived macrophages used in this study come from a mixed strain (though predominantly C57BL/6) of mice harboring a conditional *Arpc2* allele ^[66]^. This allele consists of LoxP sites flanking exon 8 of the gene encoding the p34/Arpc2 subunit of the Arp2/3 complex. In addition, these cells harbor *Ink4a-/-; Arf-/-* alleles. It is this double knockout that allows these macrophages to persist in culture in a semi-immortalized state. This is a defined genetic event rather than a spontaneously arising oncogenic immortalization. Finally, these cells also harbor a Rosa26-CreER transgenic allele that allows for conditional deletion of *Arpc2* exon 8 in the presence of tamoxifen. None of the cells used in the present study were exposed to tamoxifen, so they were considered functionally wild-type for the duration of their time in culture. The GFP-myosin IIA macrophages come from homozygous knock-in mice in the C57Bl/6 background, kindly provided by Robert Adelstein (National Heart, Lung and Blood Institute; Bethesda, MD) ^[67]^.

### Reagents

#### Antibodies

α6 integrin antibody (GoH3, ThermoFischer, #14-0495-82); p-Cofilin Ser3 antibody (77G2, Cell Signaling, #3313); p-MLC Ser20 (for western blot) (AWBMyl9F6 (F-6), ThermoFischer, #MA5-27983); GAPDH antibody (clone 6C5, ThermoFisher, #AM4300); Myosin IIA antibody (Polyclonal, Cell Signaling, #3403); Phospho-Myosin Light Chain 2 antibody (for immunofluorescence) (Polyclonal, Cell Signaling, #3674); Goat anti-Rabbit IgG Rhodamine Red-X (RRX) secondary antibody (Polyclonal, Jackson Immunoresearch, 111-295-144); Goat anti-Mouse IgG RRX secondary antibody (Polyclonal, Jackson Immunoresearch, 115-295-166); Goat anti-Rabbit IgG Alexa 488 secondary antibody (Polyclonal, Jackson Immunoresearch, 111-545-144); Goat anti-Mouse IgG Alexa 488 secondary antibody (Polyclonal, Jackson Immunoresearch, 115-545-166); Goat anti-Rat IgG RRX secondary antibody (Polyclonal, Jackson Immunoresearch, 112-295-167); Goat anti-Rat IgG Alexa 488 secondary antibody (Polyclonal, Jackson Immunoresearch, 112-545-003); Goat anti-mouse IgG, HRP-conjugated (Jackson Immunoresearch, 115-035-146); Goat anti-rabbit IgG, HRP-conjugated (Jackson immunoresearch, 111-035-144)

#### Extracellular matrix components

Poly-L-lysine (Sigma-Aldrich, #P8920); Rat tail collagen, type I (ThermoFisher, #A1048301); Fibronectin, human plasma (ThermoFisher, #33016015); Laminin 111, mouse (ThermoFisher, #23017015); Vitronectin, human plasma (Sigma-Aldrich, #5051); Rhodamine-conjugated fibronectin (Cytoskeleton Inc., FNR01-A); HiLite 488-labeled laminin (Cytoskeleton Inc., LMN02-A)

#### Other reagents

CellTracker™ Green CMFDA Dye (Invitrogen, #C7025); CellBrite® Orange: Ex/Em 549/565 nm (Biotium, #30022); Alexa Fluor™ 647 Phalloidin (Invitrogen, #A22287); Alexa Fluor™ 488 Phalloidin (Invitrogen, #A12379)

### Inhibitors and Inhibitor Use

S-nitro blebbistatin (at 7.5, 15, and 30 µM; Fisher NC0664123), Y-27632 (at 5, 10 and 20 µM; abcam ab120129), cilengitide (at 20, 40, and 60 µM; Sigma-Aldrich SML 1594-5MG). Inhibitor dilutions were made up in macrophage media. Cells were exposed to inhibitor for at least one hour prior to initiation of experiment, and inhibitor was present throughout the duration of the experiment unless otherwise noted.

### Immunofluorescence

12 mm glass coverslips were coated inside single wells of 24-well dishes, and coated with 400 µL of ECM diluted to desired concentration in sterile 1X PBS. Coverslips were incubated for 1h at 37°C. After 1 hour, coverslips were washed 3 times with sterile 1X PBS. After final wash, 400 µL of macrophage media was added to each well. Each coated coverslip was seeded with 7,500-10,000 macrophages, depending on the experiment. Macrophages were allowed to equilibrate and spread overnight in the incubator. Media was gently aspirated off the following day, leaving cells bound to the coverslip. Cells were washed once with RT non-sterile 1x PBS (hereafter referred to simply as ‘PBS’). Cold or RT (depending on antigen) 4% paraformaldehyde in Krebs Buffer or PBS (depending on antigen) was used to fix cells; incubation was 10 minutes at room temperature. Wells were then washed 3X 10 minutes with 2 mL PBS. PBS was aspirated and cells were permeabilized for 5 minutes with 0.1% Triton X-100 in PBS. Coverslips were again washed rapidly 3 times with PBS. Coverslips were then incubated for 30 minutes at room temperature with a blocking solution of 5% Bovine Serum Albumin and 5% normal goat serum in PBS and then aspirated. Primary antibody was diluted into 1% BSA in PBS (range of 1:100-1:250, depending on antibody). Coverslips were then incubated for 1 hour at room temperature. Coverslips were then washed 2x 5 minutes in RT PBS. Fluorescent secondary antibodies and phalloidin were diluted 1:500 into 1% BSA in PBS and incubated for 30 minutes at room temperature. Secondary antibody solution was then aspirated and wells were washed with 1:10,000 Hoechst diluted in PBS for 5 minutes at RT. The wells were washed once more with PBS for 5 minutes at RT. After the final 1x PBS wash, 2 uL of fluoromount G was added to a microscope slide. Using fine forceps, each coverslip was transferred cell-side-down to the fluoromount G. After a 10 minute incubation at room temperature in the dark, the coverslips were sealed with nail polish. Samples were then imaged on an Olympus IX83 epifluorescence microscope, a Zeiss 700 confocal microscope, or a Zeiss 980 Airyscan (see details, below).

### Time lapse video microscopy

Collagen, fibronectin, laminin, vitronectin and poly-L-lysine were diluted to the desired concentration in sterile 1X PBS. Glass bottomed 8-well chamber slides were coated with 300 µL of the appropriate working dilution of ECM. Chamber slides were then incubated at 37°C for 1 hour. ECM solution was then aspirated and surfaces washed rapidly 3 times with sterile 1X PBS. Macrophage media was added to each chamber, and the chamber was stored in the incubator until cells were ready to be added. <u>Labeling with tracker dye</u>: A 1:10,000 dilution of green cell tracker dye in PBS, or 1:1,000 dilution of CellBrite Cytoplasmic Membrane Dye Orange (Biotium #30022) was made in macrophage media. Macrophage media was aspirated from the culture dish and working solution of green cell tracker dye was added at room temperature for 5 minutes, and then handled according to standard macrophage passaging protocol. If orange cell tracker dye was used, the working solution was added to culture dish for 1 hour, and then cells were handled according to standard macrophage passaging protocol. <u>Seeding macrophages in chambers</u>: After labeling, 5,000 cells were added to each well of the culture chamber. The chamber slide was moved to the IX83 Olympus microscope and loaded into a Tokai Hit stage-top environmental chamber set to keep the cells at 5% CO_2_ and 37°C (see details below). Cells were allowed to equilibrate for at least 2 hours prior to imaging. Cells were then imaged at 20x magnification via relief contrast and the appropriate channel for the chosen tracker dye. In a typical experiment seven positions in each condition were chosen. The time lapse was set to image every 10 minutes for at least 16 hours. <u>Image analysis</u> to obtain migration velocity and persistence was performed in FIJI. The TrackMate plugin was used to generate track information, which was subsequently loaded into the Chemotaxis plugin to generate velocity and persistence values. The only exception to this is cells treated with blebbistatin alone, which were not fluorescently labeled due to blebbistatin’s cytotoxicity under blue light illumination as these experiments occurred before our adoption of orange tracker dye. These experiments were tracked by hand using the FIJI manual tracking plugin rather than TrackMate, but subsequent analysis with the Chemotaxis tool occurred identically to the procedure outlined above.

### Adhesion assays

Glass surfaces of multi-well chamber slides were coated with 10 µg/mL FN or 10 µg/mL LAM for 1 hour at 37°C. Chambers were then washed 3x with sterile 1X PBS and stored overnight at 4°C. PBS was aspirated the next morning at 300 µL macrophage media was added, and chamber was allowed to equilibrate in a tissue culture incubator for at least 1 hour prior to addition of cells. <u>Inhibition with α6 blocking antibody</u>: 20,000 macrophages were centrifuged for 4 minutes at 1,000x g. Excess media was aspirated and cells were re-suspended in 15 µL of 2% FBS containing a 1:1,000 dilution of orange cell tracker dye, and either α6 blocking antibody or isotype normal IgG control. Macrophages were resuspended and incubated for 37°C for 60 minutes, and were vortexed once halfway through incubation. Macrophages were centrifuged again for 4 minutes at 1,000x g and resuspended in 10 µL of macrophage media. 5 µL of this suspension (∼10,000 cells) was added to previously coated chamber wells. <u>Cilengitide treatment</u>: Cells were treated according to standard passaging protocol. 10,000 cells were plated directly into chamber wells containing Cilengitide or an equal volume of DMSO. <u>Imaging and analysis</u>: After either approach, the chamber slide was transferred to the BioTEK Cytation microscope. In each case cells were allowed to adhere and spread for 2 hours. Images were collected at this point, representing the ‘pre-wash’ population. Macrophage media was then aspirated and 2 washes with 1X PBS were used to gently wash cells on the cover glass with a P1000 pipette. The final wash was aspirated and 300 µL of macrophage media was added back, and another round of imaging was conducted to capture post-wash macrophages. Cell number was quantified from the TRITC channel (to detect cell tracker label) before and after washes to obtain the ratio of macrophages before and after washing.

### Quantitative Image Analysis

<u>Live cell velocity and persistence tracking</u>: Image files were opened with the FIJI Bio-Formats Importer plugin. An FFT bandpass filter was used to enhance contrast between cells and the background, and applied with default settings. The FIJI TrackMate file was used to identify and track fluorescently labeled cells. LoG detector was selected, and estimated ‘spot’ diameter was set such that as many artifacts as possible were excluded and as many cells as possible were detected. Simple LAP tracker was selected. Linking max distance and gap-closing distance were both set to 70.0. Gap-closing max frame gap was set to 10. Tracks were filtered based on track duration, track mean speed, and track displacement to filter out dead cells and artifacts (which typically had a speed and displacement of 0) and to avoid sampling tracks with overly short durations. Raw data was saved as .csv files and imported into the Ibidi Chemotaxis tool FIJI plugin to obtain velocity and persistence values for each cell track. <u>Cell area, elongation, and F-actin</u> <u>fluorescence</u>: The Olympus CellSens software was used to quantify cell area, elongation and fluorescence intensity from Phalloidin staining (which labels F-actin throughout the cell). Parameters were set such that small fragments of cells were excluded from batch analysis, and signals arising from multiple cells (e.g. cells touching in an image) were manually excluded from analysis. <u>Circularity quantification</u>: The freehand selection tool in FIJI was used to trace cells in relief contrast images. Circularity ranges from a value of 1 (perfectly circular) to 0 and was determined using the ‘shape descriptors’ function. <u>Internalization of Rhodamine-fibronectin</u>: Images were imported into FIJI with autoscale turned off. Display range is manually set to 0 minimum and 10,000 maximum. These values were kept consistent for images with and without Rhodamine-fibronectin. ‘Set measurements’ was used to measure area, mean gray value, and min and max gray value. Cells were traced in the relief contrast channel, then the channel selector was toggled to the fluorescent channel, where the measurements were taken. <u>Fluorescent ECM coating</u>: Images were imported into FIJI with autoscale turned off. Display range was manually set to 0 minimum and 10,000 maximum value. “Set measurements” was used to determine mean gray value, and min and max gray value. A square size was manually set on the image in the top left corner of each image, then measurement was taken. The square was then moved to the top right corner, the center, bottom left and bottom right corner, with measurement taken at every position. This was repeated for every concentration condition. <u>Adhesion assay</u>: Macrophages labeled with orange cell tracker dye were automatically detected by the BioTEK Cytation software. Cell counts were taken immediately before and after adherent cells were washed with PBS. Measurements were taken in approximately the same area. The reported value is a simple ratio between the cell count at the end of the assay divided by cell count at the start. <u>Cell shape change</u> <u>quantification</u>: A 5-hour window from each overnight time lapse experiment was used for the analysis. The same timeframes were used from each position within an experimental set. The 5-hour time lapse session was imported into FIJI. Fluorescent images of tracker dye-labeled cells were processed so that brightness and contrast were high, and fluorescent background was minimized. This allowed the entirety of the cell to be fluorescent label-positive, so that the FIJI magic wand tool could be employed to mark each cell at every time point in a semi-automated fashion. Circularity measurements for each cell at each time point were gathered in FIJI. At this point the absolute value of t_n_-t_n+1_ was taken, where n = frame number. The average shape change for each analyzed cell was then calculated. The mean of the entire cell population average was then quantified. Blebbistatin-treated cells were not fluorescently labeled, due to blebbistatin’s known cytotoxicity under blue light illumination. The analysis process for these samples was largely the same, except that the relief contrast channel was used for analysis and cell boundaries were outlined by hand. All subsequent analysis was done identically to the procedure outlined above. <u>Slingshot behavior</u> <u>quantification</u>: A single frame cell shape change value of 0.4 or higher during the 5 hour analysis window was defined as a ‘slingshot event’. This is based on empirical observation of slingshot events in comparison to known cell shape change values calculated during the event. Every cell shape change value for every cell in a population (e.g. WT cells on FN) was sorted into one dataset. The number of values higher than 0.4 was recorded and divided by the number of cells in the population. This single value was reported for each population in each experiment conducted. <u>Myosin II organization</u>: Macrophages on fibronectin or laminin were fixed with 4% paraformaldehyde and processed for microscopy as indicated above. Myosin IIA and p-MLC antibodies were used for analysis in this experiment. Individual confocal slices were hand outlined in FIJI to capture all myosin II or p-MLC signal from each slice of the z-series, and integrated pixel density was quantified. ‘Ventral’ myosin II was defined as the pool of myosin in the cell’s lowest z-frame (i.e. the side of the cell closest to the coverslip). Myosin II or p-MLC localizing to the bottom of the cell was expressed as a fraction of the total intensity across all z-slices of the cell. A larger value indicates that more of the myosin II or p-MLC signal localizes to the cell bottom.

### Microscopes

<u>Epifluorescence</u> of fixed images was done at room temperature on an Olympus IX83 outfitted with an X-cite 120 LED Boost light source (Excelitas Tech.) for fluorescent imaging. A Hammamatsu digital camera (C13440-20CU) was used to capture images. 10x, 20x or 100x objectives were used during epifluorescence experiments, depending on experimental needs. <u>Time lapse video microscopy</u> was conducted on either the Olympus IX83 or BioTEK Cytation, as noted previously in the methods. For the IX83, an INU incubation system controller from Tokai Hit was used to maintain cells in a stable environment at 37°C, 90% humidity, and 5% CO_2_. A 20X objective was employed during time lapse imaging. A BioTEK Cytation 5 was used primarily for adhesion assays. This setup includes a wide field of view camera, high performance imaging controller and a laser autofocus add-on. Imaging can be done at 4X, 10X or 20X magnification. <u>Confocal</u>: A Zeiss LSM 700 confocal microscope housed at the USUHS Biological Instrumentation Center (BIC) was employed for higher resolution imaging of fixed samples using a 40X oil immersion objective. A Zeiss 980 Airyscan housed at the USUHS BIC was employed for GFP-myosin IIA imaging using a 63X oil immersion objective. All confocal and Airyscan imaging was done with fluorescent filter sets appropriate for each fluorophore.

### Western blotting

<u>Protein lysis</u>: Macrophages were washed 3x with ice-cold PBS. After final PBS aspiration, cells were kept on ice and RIPA buffer (Sigma R0278-50ML) with 1X protease inhibitor (Thermo A32953) and 1X phosphatase inhibitor (Sigma 4906845001) was added to cells. Cells were then scraped thoroughly and transferred to Eppendorf tubes, which were kept on ice for 15 minutes. Tubes were then centrifuged at 4°C in a pre-chilled microcentrifuge for 10 minutes. The supernatant was transferred to a new tube and the pellet was discarded. Precision Red (Cytoskeleton ADV02-A) was used to quantify protein amount in RIPA lysate. Analysis was done via plate reader. Samples were prepared based on protein reading, and typically 8 µg of protein sample was loaded per well for electrophoresis. <u>Electrophoresis</u>: Samples were combined with 1X Bolt LDS Sample Buffer (ThermoFisher B0007) and 1X NuPAGE Sample Reducing Agent (Novex NP0009), and remaining volume was made up with excess RIPA buffer to ensure that both an equal volume and protein concentration was maintained across samples. Samples were vortexed briefly, heated to 70°C for 10 minutes, and centrifuged at max speed for 1 minute. Samples were then loaded onto a Bolt 4-12% Protein Gel (ThermoFisher NW04125BOX). The gel electrophoresis chamber was filled with 1X NuPAGE MES SDS Running Buffer (Thermo NP0002) and run at 200 V. Electrophoresis was stopped when the loading dye front reached the bottom of the gel. <u>Transfer</u>: A traditional wet transfer to PVDF membrane (Bio-Rad 1620174) was done for 1 hour on ice with chilled transfer buffer using 0.35 A (constant amps setting). Transfer buffer was diluted to 1X from 10X stock (for 1L of 10X transfer buffer: 19.70 g Trizma HCl, 15.14 g Trizma base, 142.63 g glycine, adjusted to pH 8.3). After dilution methanol was added to a final concentration of 20%. <u>Blotting</u>: Membranes were placed directly in 5% milk (in 1X TBST) for 1 hour at room temperature. Membranes were then washed, and primary antibody diluted in 5% BSA (in 1X TBST) or 5% milk (in 1X TBST) was added overnight in the cold room with gentle agitation. Membranes were then washed 3x 5 minutes with 1X TBST. The membrane was then incubated for 30 minutes with 5% milk in TBST containing a 1:10,000 dilution of HRP-conjugated secondary antibody. The blot was then washed 3×5 minutes with 1X TBST. HRP signal was activated with SuperSignal West Pico Chemiluminescent Substrate and imaged on an Amersham Imager 680. FIJI was used to analyze band intensity, using identically sized boxes to determine total pixel intensity in each sample for each protein of interest. All intensities were normalized to GAPDH to control for differences in loading across samples. Uncropped blots can be found in Supplemental Figure S5. Images were evaluated to ensure that only un-saturated exposures were analyzed. Any brightness and contrast changes were applied equally to the entire image.

### Statistical analysis

The Kruskal-Wallis with Dunn multiple comparisons test was used to assess significance in experiments where a normal distribution of the dataset could not be assumed. When only 2 experimental conditions were tested, we used Mann-Whitney when we could not assume normality. Unpaired t-tests and ANOVAs were used when normality tests indicated a normal distribution of the data. All statistics were calculated using GraphPad Prism, and significance was assumed if p ≤ 0.05. More information on each statistical test can be found in the relevant figure legend panel.

## LIST OF SUPPLEMENTARY MATERIALS

**Figure S1**: Additional p-values for relationships between cells plated on various ECMs encompassing Figures 1B-D and 1E-F.

**Figure S2**: Additional data related to main Figures 2

**Figure S3**: Additional data related to main Figures 4 and 6

**Figure S4**: Uncropped images of main Figure data

**Figure S5**: Uncropped blots of main Figure data

**Supplemental Movies 1:** WT macrophages, random migration on 10 µg/mL fibronectin

**Supplemental Movie 2**: WT macrophages, random migration on 10 µg/mL lamini

**Supplemental Movie 3**: WT macrophages, random migration on 1 µg/mL fibronectin

**Supplemental Movie 4**: WT macrophages, random migration on 1 µg/mL laminin

**Supplemental Movie 5**: WT macrophages, random migration on 10 µg/mL laminin + 30 µM blebbistatin

**Supplemental Movie 6**: WT macrophages, random migration on 10 µg/mL laminin + 20 µM Y-27632 and 15 µM blebbistatin

**Supplemental Movie 7**: WT macrophages, random migration on 10 µg/mL fibronectin + 20 µM Y-27632 and 15 µM blebbistatin

**Supplemental Movie 8**: WT macrophages, random migration on 10 ug/mL fibronectin + 10 ug/mL laminin

**Supplemental Movie 9**: WT macrophages, random migration on 10 ug/mL fibronectin + 30 ug/mL laminin

**Supplemental Movie 10**: WT macrophages, random migration on 30 ug/mL fibronectin + 10 ug/mL laminin

**Supplemental Movie 11**: WT macrophages, random migration on 10 ug/mL laminin + 60 µM cilengitide

**Supplemental Movie 12**: WT macrophages, random migration 10 ug/mL laminin + 40 µM cilengitide + 20 µM Y-27632

**Supplemental Movie 13**: WT macrophages, random migration 10 ug/mL laminin + 40 µM cilengitide + 15 µM blebbistatin

## ACKNOWLEDGEMENTS

We thank Dennis McDaniel and the USUHS Biomedical Instrumentation Center for help with confocal microscopy. GFP-myosin IIA knock-in macrophages were provided by Dillon Schrock and John Hammer, and were a kind gift from Robert Adelstein (National Heart, Lung, and Blood Institute; Bethesda, MD).

## AUTHOR CONTRIBUTIONS

M.S. planned and performed experiments, analyzed data, and wrote and edited the manuscript. A.L performed experiments and analyzed data. J.R. supervised the project, planned and performed experiments, wrote and edited the manuscript, and secured funding. All authors had the opportunity to review and comment on the manuscript prior to submission.

## DATA AVAILABILITY STATEMENT

All primary data will be openly available.

---

**Supplemental Figure 1:**
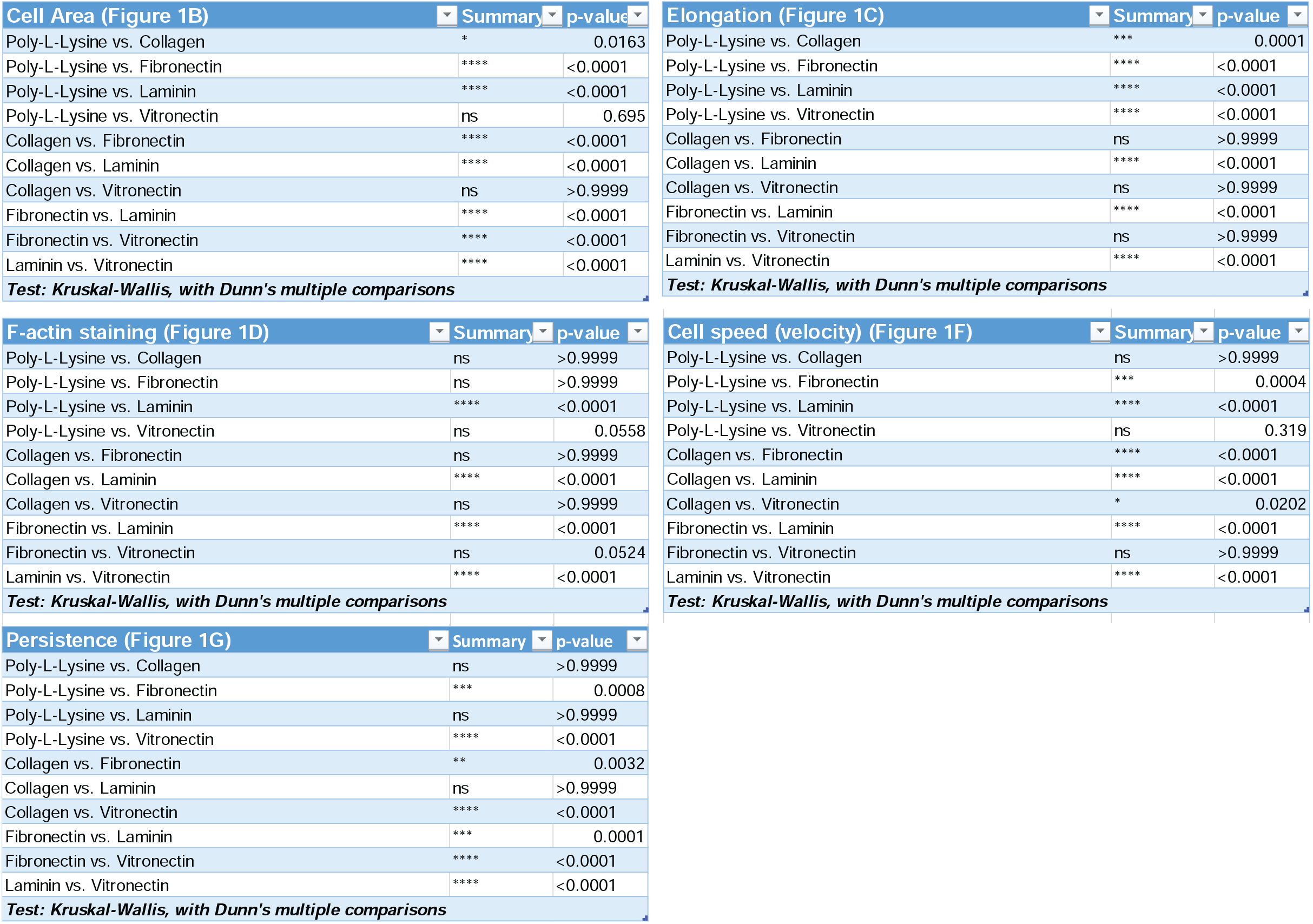
Additional experiments related to main Figure 1. Complete list of statistical relationships between measured values for cells plated on one ECM versus another. More information about these experiments can be found in the main figure legend indicated for each grouping.

**Supplemental Figure 2:**
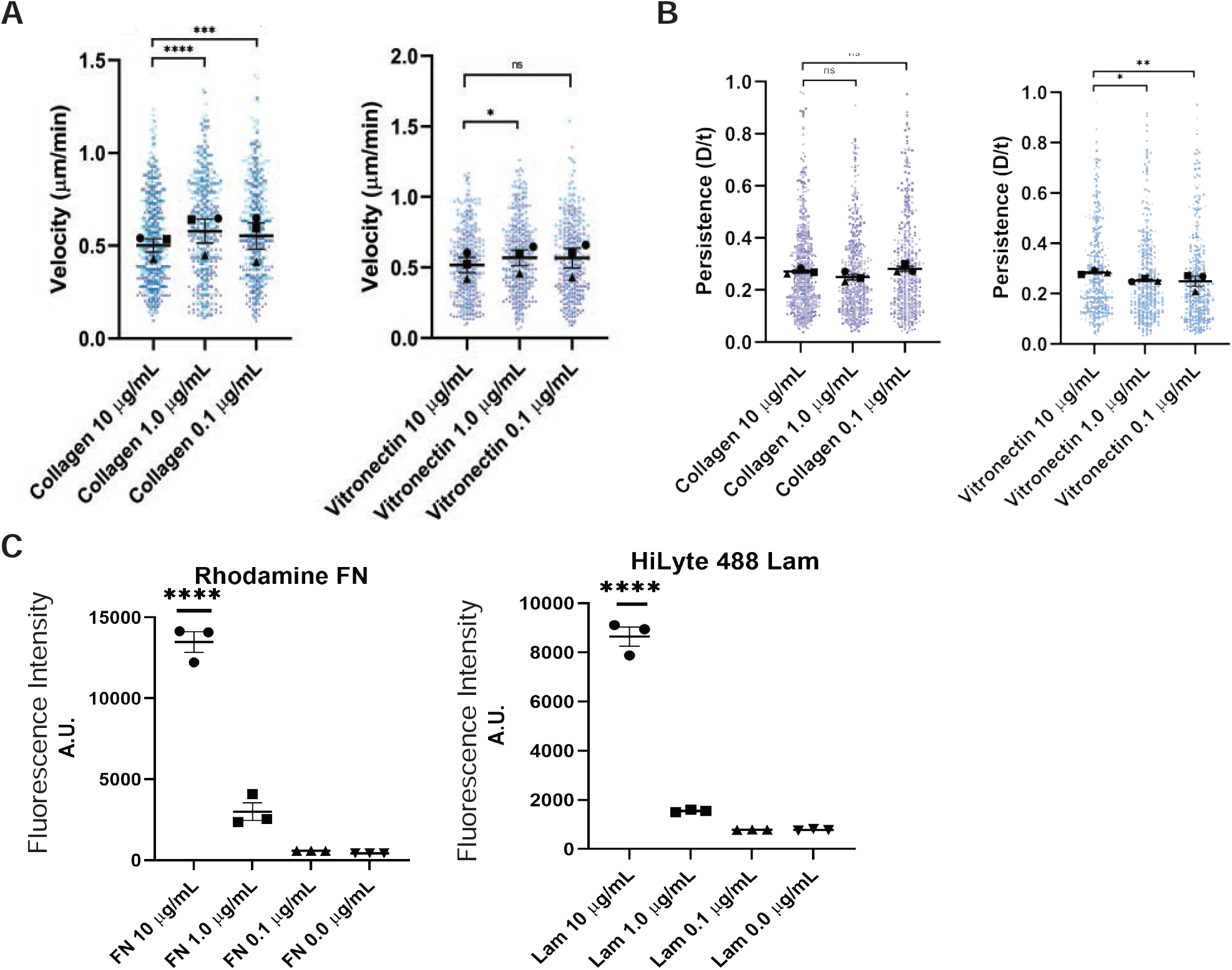
Additional experiments related to main Figure 2. (A) Velocity (cell speed) in microns/minute. (**** = p<0.0001, *** = p=0.0003, * = p=0.0149, ns=not significant. Col 10 μg/mL n=543 tracks, Col 1.0 μg/mL n=459 tracks, Col 0.1 μg/mL n=438 tracks, VN 10 μg/mL n=365 tracks, VN 1.0 μg/mL n=402 tracks, VN 0.1 μg/mL n=378 tracks pooled from 3 experiments). Means and standard error of the mean for each experiment are represented by black symbols. All data points are plotted and symbol shape corresponds to the trial mean. Statistical analysis was Kruskal-Wallis with Dunn multiple comparisons test. (B) Persistence (d/T) (**p = 0.0016, *p = 0.0261, ns = not significant. Track n is the same as reported in panel A. Means and standard error of the mean for each experiment are represented by black symbols. All data points are plotted and symbol shape corresponds to the trial mean. Statistical analysis was Kruskal-Wallis with Dunn multiple comparisons test. (C) Mean fluorescence intensity of Rhodamine fibronectin (FN) and HiLyte 488 laminin (LAM)-labeled coverslips treated with the indicated concentrations. Fluorescence images were collected from 5 zones and averaged for 3 fields of view and averaged. Error bars represent standard error of the mean. ****p < 0.0001. Statistical significance was assessed with ordinary one-way ANOVA with Tukey’s multiple comparisons test.

**Supplemental Figure 3:**
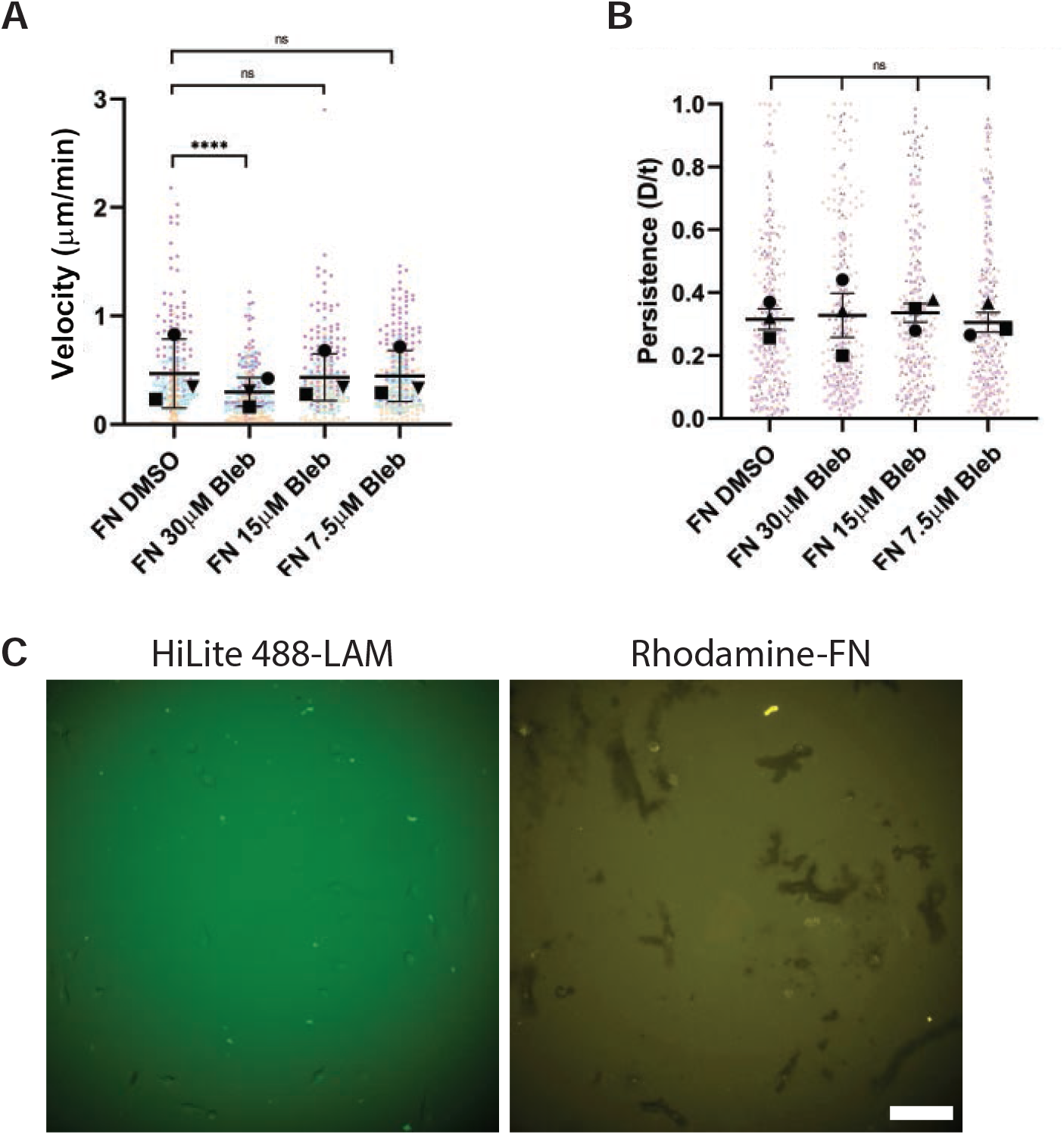
Additional experiments related to main Figures 4 and 6. (A) Velocity (Cell speed) and (B) Persistence (d/T) for macrophages randomly migrating overnight in the presence of the indicated dose of blebbistatin on 10 µg/mL fibronectin. Black symbols represent experimental means, bars are standard error of the mean. Every experimental data point is plotted and color-coded based on experiment. n = at least 248 tracks per condition, pooled from three independent experiments. ***p < 0.0001. Statistical significance was assessed using Kruskal-Wallis and Dunn’s multiple comparisons test. (C) Example images demonstrating simultaneous deposition of fluorescent laminin and fluorescent fibronectin on a single substrate. These fields of view are uncropped images of the same fields presented in Figure 6C in the ‘Rhodamine-Fibronectin’ condition. Scale bar = 100 microns.

**Supplemental Figure 4:**
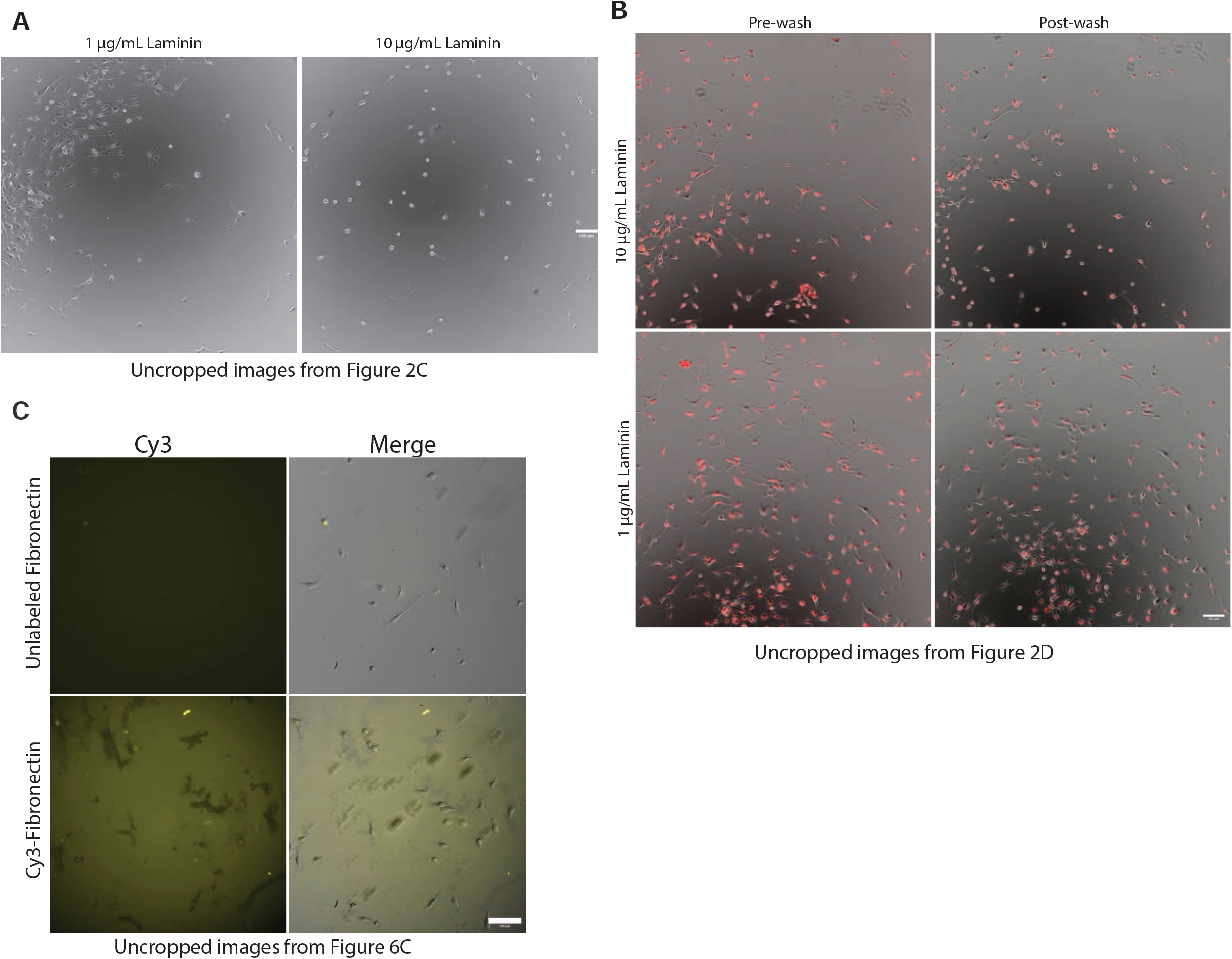
Uncropped versions of images used in main figures. More information regarding these images can be found in the indicated main figure panel. All scale bars in this Supplemental Figure are the same size as the bars in the corresponding main figure panel.

**Supplemental Figure 5:**
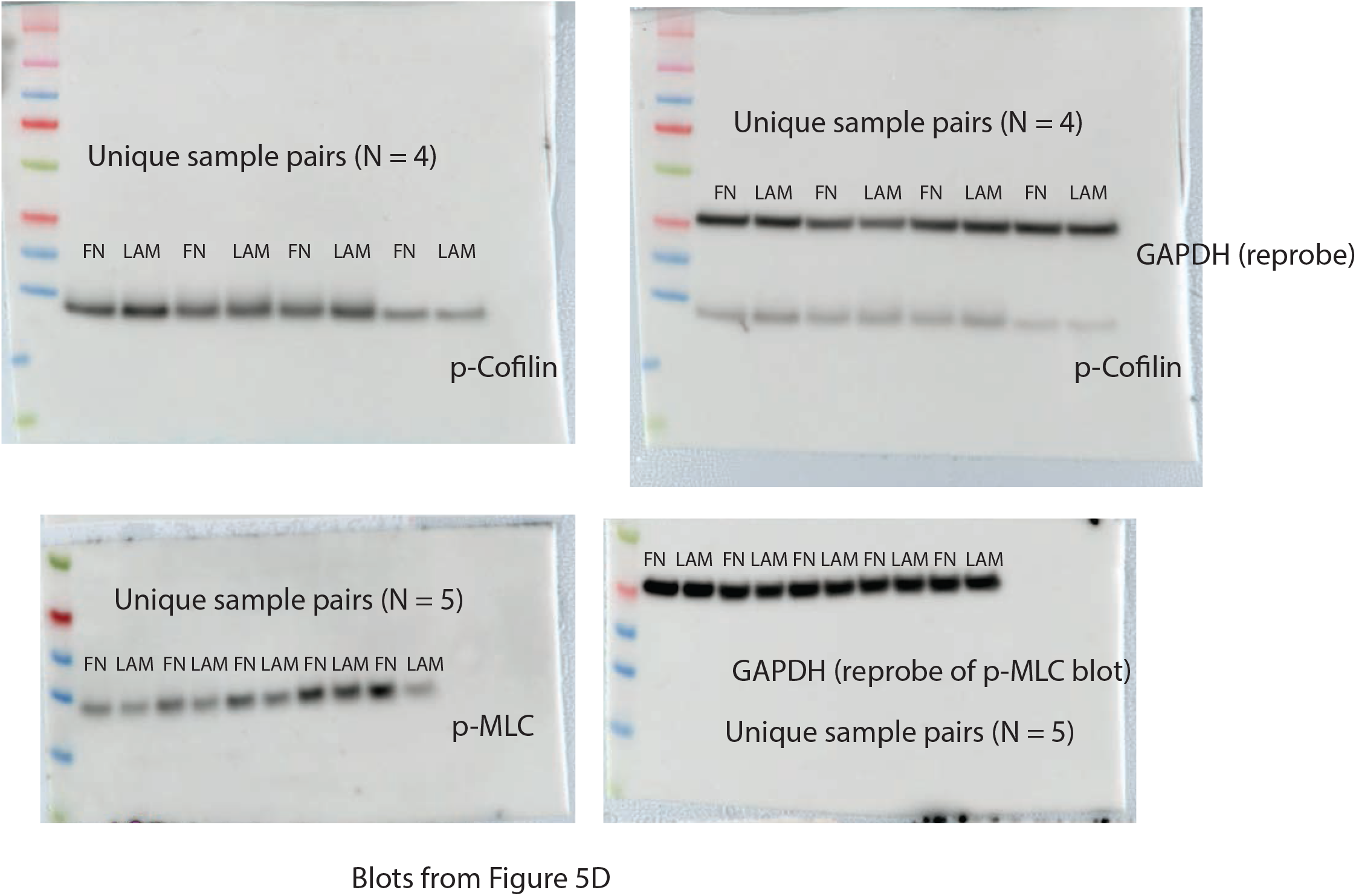
Uncropped blots. Each blot has been reproduced here demonstrating all of the biological replicates (N = 4 or 5) used in the quantitative analysis in Figure 5D. Each blot has been labeled with the protein that was probed. The blots have been merged with its corresponding pvdf membrane, which also includes the protein molecular weight marker. As the GAPDH antibody was raised in mouse (and the others in rabbit), GAPDH was directly reprobed after initial probing with p-MLC or p-cofilin antibody. These blots were not stripped prior incubation with the GAPDH antibody.

